# Simultaneous isolation of proximal and distal lung progenitor cells from individual mice using a 3D printed guide reduces proximal cell contamination of distal lung epithelial cell isolations

**DOI:** 10.1101/2022.05.10.491312

**Authors:** Hani N. Alsafadi, John Stegmayr, Victoria Ptasinski, Iran Silva, Margareta Mittendorfer, Lynne Murray, Darcy E. Wagner

## Abstract

The respiratory epithelium consists of multiple, functionally distinct cell-types and is maintained by regionally-specific progenitor populations which repair the epithelium following injury. Several *in vitro* methods exist for studying lung epithelial repair using primary murine lung epithelial cells, but isolation methods are hampered by a lack of surface markers distinguishing epithelial progenitors along the respiratory epithelium. Here, we developed a 3D-printed lobe divider (3DLD) to aid in simultaneous isolation of proximal versus distal lung epithelial progenitors from individual mice which give rise to differentiated epithelia in multiple *in vitro* assays. In contrast to 3DLD-isolated distal progenitor cells, classic manual tracheal ligation methods followed by lobe removal resulted in co-isolation of rare proximal progenitors with distal cells which altered the transcriptional landscape of distal organoid cultures. Thus, cell isolation with the 3DLD generates reproducible distal versus proximal progenitor populations and minimizes the potential for contaminating populations to confound *in vitro* assays.

**Highlights:**

- 3DLD reproducibly separates lung lobes and extrapulmonary airways (bronchi/trachea)
- 3DLD cell isolation yields consistent isolation of distal epithelial cells (DECs)
- Contamination of proximal cells in classic DEC isolations may alter *in vitro* results
- 3DLD allows for simultaneous isolation of proximal and DECs from single animals

**eTOC blurb:** Alsafadi et al. describes a new method for simultaneous isolation of lung epithelial proximal and distal progenitors using the aid of a 3D printed device (3DLD). Both isolated cell types differentiate in multiple *in vitro* assays. The 3DLD guide minimized contamination of proximal cells in distal cell isolations whose presence can alter the transcriptional landscape of distal epithelial organoids.

## Introduction

The respiratory epithelium is a complex structure consisting of several distinct cell types distributed within the nasal and trachea passages leading to the branching airways and terminating in the distal alveoli where gas exchange occurs (Rock et al., 2010; Volckaert and De Langhe, 2014). Although the lung is known to have a limited regenerative capacity, it contains several regional-specific progenitors which have been shown to be capable of proliferation and subsequent differentiation to repair damaged lung epithelium (Chen et al., 2012). The cell types responsible for epithelial repair and their origin has been described in several seminal papers describing their location within the lung as well as their regenerative capacity following specific injuries (Barkauskas et al., 2013; Kim et al., 2005; McQualter et al., 2010; Rawlins et al., 2009; Reynolds et al., 2000; Rock et al., 2009).

The tracheal epithelium is maintained by a heterogenous pool of basal cells (Rock *et al*., 2009) while the normal bronchioles and alveolar epithelium are maintained by Scgba1+ cells (termed club cells) and Sftpc+ alveolar type II (ATII) cells, the progenitor cell of the alveolus (Barkauskas *et al*., 2013; Rawlins *et al*., 2009). Additionally, subpopulations such as Itga6+ Itgb4+ alveolar progenitors and bi-potent progenitor populations such as bronchioalveolar stem cells (BASCs) (Scgb1a+ Sftpc+) and Hopx+ have also been shown to be capable of giving rise to both small airway and alveolar cell types, respectively (Chapman et al., 2011; Jain et al., 2015; Kim *et al*., 2005; Rawlins *et al*., 2009). Single cell RNA sequencing (scRNAseq) studies are continuously improving our understanding of the diverse cellular landscape in the lung, including progenitor populations which arise and expand during disease and especially the presence of multiple transitory populations in disease in both human lung tissue and during murine models of diseases (Sikkema et al., 2022). These transitional states are defined by the co-expression of multiple phenotypic markers associated with previously described adult cell types and once thought of to be mutually exclusive in defining cell populations. Therefore, hereafter, for simplicity, we will generally refer to the basal epithelial cell populations located in murine tracheas as proximal progenitors and those from parenchymal cell isolations as distal progenitors.

Several different *in vitro* and *in vivo* approaches have been used to identify the cell types involved in lung repair and regeneration. *In vitro* studies of isolated cells on tissue culture plastic or floating collagen gels were initially critical in characterizing the transition of ATII to ATI cells (Danto et al., 1995). In addition, *in vivo* lineage tracing has been instrumental in identifying several of the regionally specific progenitor populations (Tata and Rajagopal, 2017). While lineage tracing has long been the gold standard for identifying progenitor populations, the promotor regions used to mark them have been found to be leaky and recent single cell studies have further confirmed that these markers are not limited to an individual cell type, even in healthy lungs and airways (Joshi et al., 2019; Montoro et al., 2018; Perl et al., 2009; Rawlins, 2008; Strunz et al., 2020). Thus, an increasing number of recent studies have utilized organoids derived from isolated cells based on the presence or absence of surface markers to predict and monitor the cell populations responsible for repair and regeneration in the lung epithelium (Barkauskas et al., 2017). However, the proportion of proliferating cells in this assay is known to be small and is highly dependent on the ability to isolate well-defined progenitor populations.

While several isolation techniques exist to isolate lung epithelial progenitors from the trachea or parenchymal tissue (Dobbs et al., 1986; Jansing et al., 2018; Lam et al., 2011; Vanderbilt et al., 2015; You et al., 2002), a protocol to simultaneously isolate murine lung epithelial progenitors from the distal epithelium (e.g. ATII cells, SPC+) and proximal epithelium (basal cells, p63+/Krt5+) has not been described due to a lack of unique surface marker(s) distinguishing the entire pool of murine proximal or distal lung progenitor cells. Furthermore, as some progenitor populations, such as Scgb1a1+ cells, reside in both the proximal and distal airways, sorting approaches using lineage tracing can be challenging (Kim *et al*., 2005; Montoro *et al*., 2018; Rawlins *et al*., 2009). Current approaches use instillation of dissociating agents through the trachea, most often followed by ligation of the trachea or, less commonly due to technical challenges, at the level of the main bronchi (Jansing *et al*., 2018; Messier et al., 2012). Single cell suspensions are then collected from the parenchymal tissue to isolate the distal epithelial cells, often through epithelial cell adhesion molecule (EpCAM) positivity or through lineage tracing approaches for isolation of specific subsets using specific markers, such as *Sftpc*. However, there is a high diversity of cell types which can become labeled, with some reports finding only 25-60% of ATII cells captured using *Sftpc* based approaches (Chen *et al*., 2012; Perl *et al*., 2009; Vanderbilt *et al*., 2015). Furthermore, *Sftpc* expression has been found to be expressed in multiple cell types of the airway epithelium in lineage tracing studies and recent scRNAseq datasets (Perl *et al*., 2009; Rawlins, 2008; Strunz *et al*., 2020), calling into question its specificity for cell isolations. On the other hand, EpCAM is commonly used for distal progenitor cell isolation, with or without additional surface markers, such as Sca1, to demarcate different regenerative subpopulations (Louie et al., 2022; McQualter *et al*., 2010). However, both proximal and distal epithelial progenitors express EpCAM and thus a small number of proximal cells may be inadvertently co-isolated in such approaches. This is especially problematic in organoid studies where around 0.5-2% of cells are identified as giving rise to organoids (Barkauskas *et al*., 2017).

Chronic lung diseases often affect both proximal and distal epithelium, but the extent and type of injury differs (Rock *et al*., 2010). Recent evidence has indicated that the response and expansion of progenitor cells are disease specific in both murine models and clinical disease (Sikkema *et al*., 2022; Vaughan et al., 2015; Zuo et al., 2015). In particular, increases in phenotypic markers traditionally associated with proximal cells has been observed in distal lung tissue for murine models of acute and chronic lung injury as well as human disease, but the origin of these cells remains incompletely understood (i.e. expansion of rare cell types residing in the distal lung, acquisition of phenotypic markers by distal lung cells, differentiation of a local progenitor cell or migration)(Vaughan et al., 2014; Zuo *et al*., 2015). Thus, in addition to reducing overall animal numbers, controlled and simultaneous isolation of both proximal and distal epithelial cells from the same mouse airway and lung epithelium based on their physical location, and not phenotypic identity, may offer opportunities to further understand their origin and role in disease.

In the current study, we develop and validate a new isolation method to simultaneously and more precisely isolate both proximal and distal progenitors from individual mice through the controlled physical separation of the trachea and lung lobes with the aid of a 3D printed device. Furthermore, we show that precise ligation of the bronchi results in isolation of a more homogenous pool of epithelial progenitor cells from the distal lung and minimizes the presence of rare contaminating proximal cells in this isolated population. In addition, we show that the use of the 3DLD does not impact cellular viability for either the proximal or distal progenitor cells and that these cells perform comparably to cells isolated using the traditional methods in multiple state-of-the-art *in vitro* assays. Finally, we show that while organoids grown from distal cells isolated using the 3DLD or classic method have a similar transcriptional trajectory, there is a subset of genes which are only upregulated in organoids grown from classically isolated distal cells and these genes are associated with cell types of the proximal airway epithelium.

## Results

### 3D printed lobe divider (3DLD) allows for efficient separation between the murine trachea, bronchi and lung lobes

Protocols for isolating proximal and distal epithelial cells differ in the way dissociation solutions are introduced to the tissue as well as the type of enzymes used. Classically described protocols to isolate epithelial progenitor cells residing in the distal lung or airways (referred to as distal progenitors in this study) requires inflation of the lung lobes with enzymatic solutions administered through the trachea (e.g. dispase) while the isolation of mouse tracheal epithelial cells (referred to as proximal progenitors in this study) is achieved by incubating the dissected trachea in an enzymatic solution (e.g. pronase) (Jansing *et al*., 2018; Lam *et al*., 2011). To achieve isolation of both cell types simultaneously, the trachea must first be used as a passage to administer the distal dissociation solution for isolation of distal progenitors, followed by quick ligation of the pulmonary lobes to free the trachea for subsequent washing away of the distal dissociation solution. While this process can be done manually, reproducible placement of surgical knots on the most distal portion of the trachea or the bronchi is challenging and may lead to rare co-isolation of basal cells if co-incubated with enzymatic dissociation solutions. Thus, we designed a 3D printed lobe divider (3DLD) device to test if this could aid in reproducible and consistent separation of the left and right lung lobes at the level of the main bronchi as well as quick dissection of the trachea (Figures 1A-C). We incorporated a locking mechanism on each side of the device to prevent backflow of the dissociation solution into the trachea and to allow time for completion of the surgical knots and rapid dissection of the trachea (Figure 1C).

**Figure 1.**
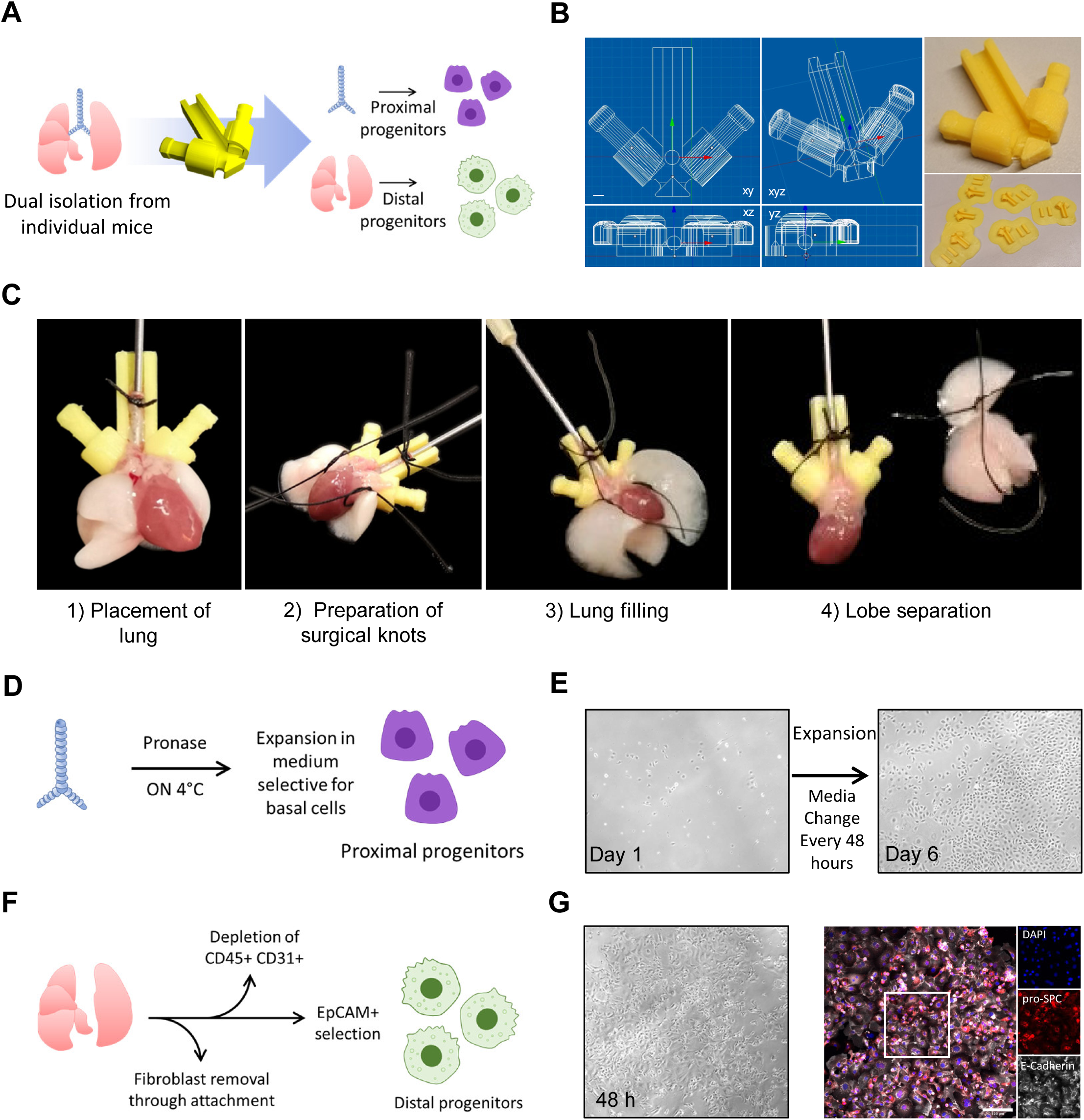
3D printed lobe divider (3DLD) allows for efficient separation between the murine lung trachea and lobes for isolating proximal and distal progenitor cells from individual mice. (A) Overview of simultaneous isolation of proximal and distal epithelial cells using 3DLD. (B) Orthogonal and isometric views of 3DLD and final printed product; (C) Overview of step-by-step 3DLD procedure; (D) Isolation and expansion of proximal progenitors; (E) representative phase contrast of proximal progenitors after 6 days of expansion. Taken with a 10X objective. (F) Overview of distal epithelial cell isolation (G) Representative immunofluorescent staining of pro-surfactant protein C (pro-SPC) and E-cadherin of distal epithelial cells (Taken with a 20X objective).

After confirming that our prototype worked for separating the lobes, we tested whether its use affected cell isolations for either distal or proximal cells using classically described techniques (Jansing *et al*., 2018; Lam *et al*., 2011) as well as in commonly used downstream assays such as *in vitro* monolayer culture (Figures 1D, 1F). As previous papers describing the isolation of proximal progenitor cells do not first expose these cells to dispase, we first confirmed that brief enzymatic exposure to dispase in the 3DLD did not negatively impact the ability of isolated proximal cells to proliferate in culture. We found that these cells readily expanded to form confluent monolayers *in vitro* after 6 days (Figure 1D, E). Similarly, we found that distal progenitor cells isolated with the 3DLD were viable and formed E-cadherin and pro-SPC positive monolayers after 48 hours, in line with previous work (Jansing *et al*., 2018) (Figure 1F, G). Thus, we found that the 3DLD allows for simultaneous isolation of progenitor cells from both the proximal and distal lung epithelial compartments from a single animal and that these cells were able to proliferate *in vitro* and retained their well-known phenotypic markers.

### Expanded proximal progenitors isolated with 3DLD differentiate into various cell-types in air-liquid interface (ALI) cultures and organoid assays

As any alterations in isolation protocols could alter the viability or composition of cell populations isolated, we next characterized the cells isolated with the 3DLD method more extensively. We first validated that the use of the 3DLD did not alter the ability of proximal progenitor cells isolated from single animals to be expanded and differentiated into mature, differentiated airway epithelial subtypes. We tested the expanded cells’ differentiation capacity using two well-established and commonly used *in vitro* assays: air-liquid interface (ALI) cultures and organoid formation (Figure 2). After 28 days of air exposure in ALI, the proximal progenitors from individual mice developed confluent monolayers and a pseudostratified and polarized epithelium with Krt5+ basal cells (Figure S1). Histological and scanning electron microscopy also confirmed morphological evidence of ciliated cells as well as the presence of alpha-tubulin positive cells restricted to the apical surface of the monolayers, confirming that the modified isolation protocol using the 3DLD does not alter the ability of the progenitor cells isolated and expanded to form differentiated and polarized monolayers (Figure S1-2 and Figure 2B). While morphological and histological assessment confirms the ability for the isolated proximal progenitors to differentiate, the choices of phenotypic markers is somewhat subjective. Thus, we performed bulk RNAseq on the proximal progenitors before and after culture at ALI to assess transcriptional changes comprehensively and objectively over time. Principle component analysis (PCA) of the full gene expression profile shows clear separation between ALI-differentiated cells and the proximal progenitors (Figure 2C), indicating major differences in the transcriptional profile between these two *in vitro* conditions. We found that 9226 genes were differentially expressed between these two populations (log fold change > 1 and p-value< 0.05; Figure 2D), several of which are known to mark various differentiated proximal epithelial cell populations (Figure 2E, Table S2).

**Figure 2.**
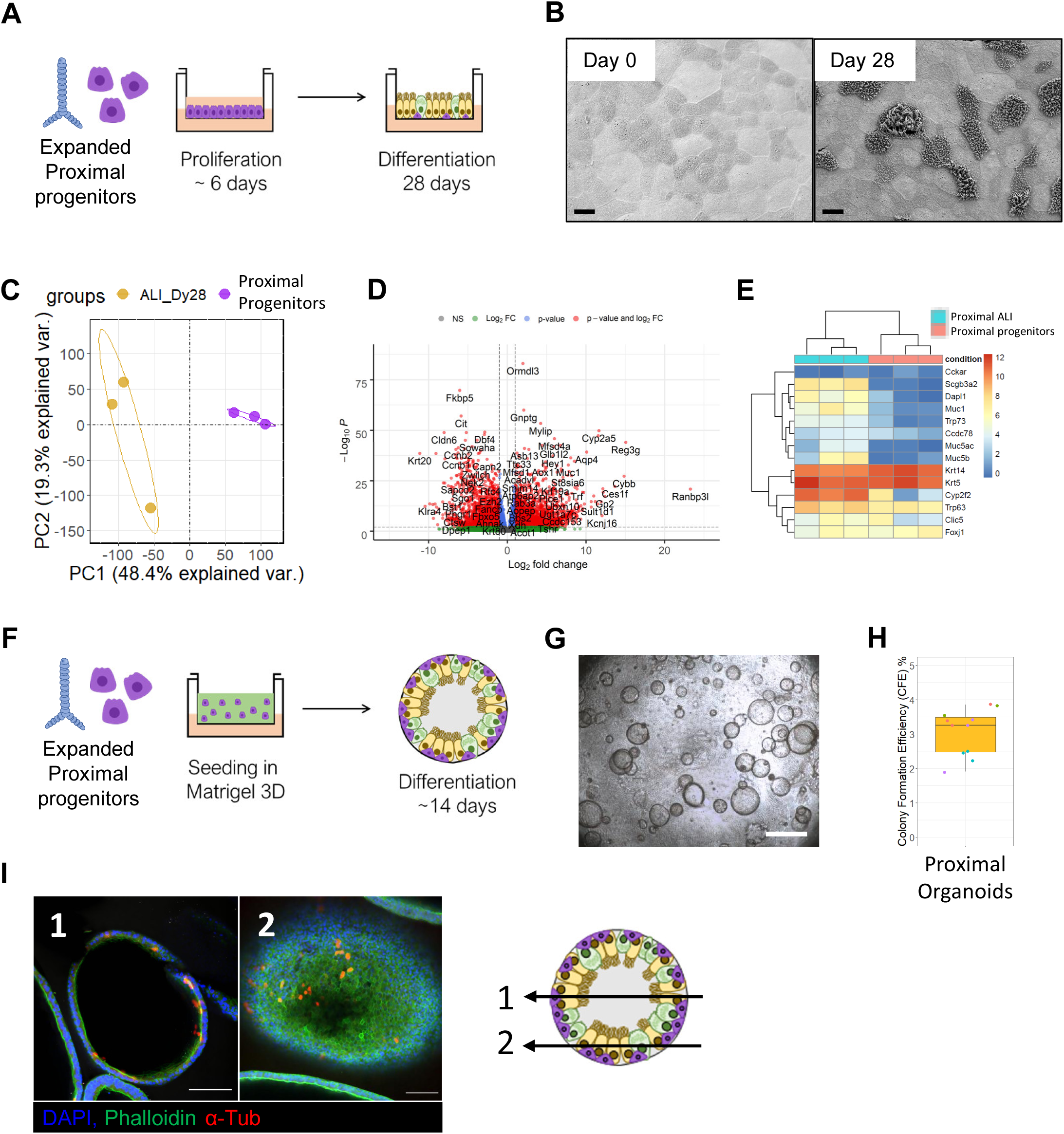
Expanded mouse proximal progenitor epithelial cells isolated with 3DLD differentiate into various cell-types in air-liquid interface (ALI) cultures and organoid assays. (A) Schematics for ALI differentiation and organoid formation assay of proximal progenitors. (B) Representative scanning electron microscopy (SEM) of ALI culture before lifting to air and 28 days after. Original magnification 1000x taken at an accelerating voltage of 3.0kV; Scalebar: 10 μm. See also Figure S1 and Figure S2. (C) Principal Component Analysis (PCA) of proximal epithelial progenitors before and after 28 days of culture in ALI. Yellow: ALI at day 28, Purple: Proximal Progenitors. (D) Volcano plot of differentially expressed genes between expanded proximal progenitors before vs after differentiation in ALI for 28 days (E) Heatmap of select phenotypic markers of differentiated proximal epithelium (F) Schematics of organoid formation assay of proximal progenitors. (G) Representative phase contrast image of differentiated proximal organoids at day 14. n=4 individual mice independently; Scalebar: 500 μm (H) Colony formation efficiency (CFE) of proximal organoids after differentiation for 14 days. n=4; 3 wells quantified per experiment represented by dots colored by mouse (I) IF staining of organoid culture after 14 days. Acetylated α-tubulin (α-tub), cytokeratin 5 (Krt5). Scalebar: 100 μm.

Next, we sought to confirm that expanded proximal progenitors isolated using the 3DLD were able to form organoids (Figure 2F-H). We found that proximal progenitors isolated using the 3DLD method formed organoids with a colony formation efficacy (CFE) of 3.1 % which is consistent with previous studies (Rock *et al*., 2009)(Fig 2H) . Furthermore, the spheroids contained alpha-tubulin positive cells, a phenotypic marker for ciliated cells, confirming differentiation in the organoid cultures (Fig 2I). Thus, we found that the alterations introduced via the 3DLD isolation procedure did not affect the ability of proximal progenitor cells to be used in at least two common *in vitro* assays.

### EpCAM+ distal progenitors differentiate into organoids with different morphologies with 3DLD isolation

In comparison to classic isolation methods for distal progenitor cells, the main alteration in the 3DLD isolation protocol is the ability to precisely remove the trachea and the extrapulmonary main left and right bronchi prior to incubation with the dissociation buffer. We first sought to compare the 3DLD isolation technique with classical protocols to evaluate potential alterations in organoid formation (Figure 3A). We found that distal progenitors from both isolation methods formed organoids with similar CFE to those reported in previous studies (Figure 3B) but the CFE was significantly higher in the 3DLD organoids (Figure 3C). Further, while both methods produced organoids of various sizes, we found the 3DLD organoids were smaller in diameter and had a narrower cross-sectional area distribution (Figure 3D-E, Figure S3). Staining of individual phenotypic markers within organoids can help yield insight into the differentiation state of cells within organoids, but it is a subjective approach. On the other hand, while scRNAseq can be used to characterize the cellular-level transcriptional landscape within distal progenitor derived organoids (Louie *et al*., 2022), it is known to be limited with regard to the sequencing depth and it is well known that some cell types, such as ATI cells, are highly sensitive to the digestion protocols and droplet-based techniques required for scRNAseq (Riemondy et al., 2019). Thus, we opted to perform bulk RNAseq from RNA isolated from freshly isolated pellets following isolation as well as the entire organoid culture well after 14 days to provide a comprehensive overview across the entire transcriptional landscape of the organoids formed using the classic or 3DLD cell isolation. Initial PCA analysis demonstrated that the two isolation methods result in highly disparate transcriptomic profiles (Figure 3F) with lower gene variance detected in cells isolated with the 3DLD (Figure S4). We found 3270 differentially expressed genes between distal progenitors isolated with the 3DLD or classic method (Figure 3G). Interestingly, among the most differentially expressed genes are those known to be associated with the proximal epithelium, such as *Krt14*, as well as genes recently associated with highly proliferative basal cells (*Pbk*)(Succony et al., 2021). However, organoids formed from both isolation methods seemed to result in transcriptional profiles which were distinct and more similar to one another than the differences we observed in the initial cell populations from each isolation (Figure 3F).

**Figure 3.**
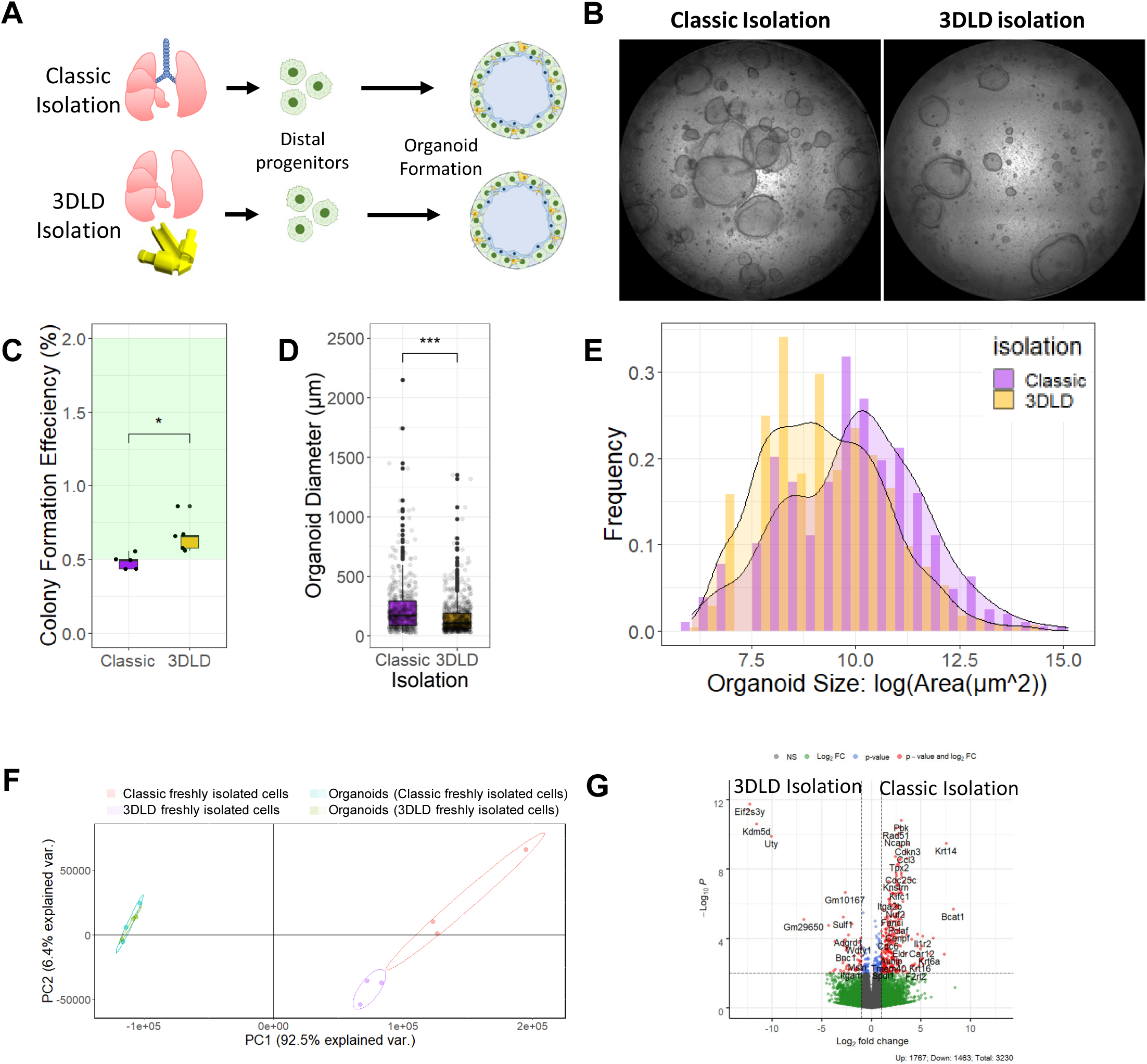
3DLD isolation alters organoid formation from EpCAM+ distal progenitors. (A) Overview of organoid formation assay using classic or 3DLD isolation methods. (B) Representative brightfield images of stitched montages of formed organoids after 14 days of culture. n=4 individual mice; Scalebar: 1mm. (C) Colony formation efficiency (CFE) from each isolation method (n=6); *p<0.05; Mann-Whitney U non-parametric t test. (D) Organoid diameter; ***p<0.0001; Mann-Whitney U non-parametric t test. (E) Frequency size distribution of distal organoids derived from distal progenitors isolated with classic or 3DLD methods. (F) Principal Component Analysis (PCA) of distal progenitors isolated with classic or 3DLD methods before and after differentiation in organoid culture. (G) Volcano plot of differentially expressed genes between the cell pellets of 3DLD and classic isolation methods before differentiation. Related to Figure S3-S4.

### Deconvolution of bulk RNAseq of distal progenitor pellets reveals higher proportions of contaminating proximal airway cells using a classic isolation protocol

To further and more comprehensively explore the transcriptional differences observed in the initial cell pellets (Figure 3F-G), we used computational deconvolution of our bulk RNAseq to give insights into predicted cellular diversity. Computational deconvolution takes advantage of the annotated cellular level information that can be derived from scRNAseq datasets to computationally dissolve the complexity within bulk RNAseq samples to predict the proportions of cell. It has been shown to accurately predict experimentally mixed cell types, and is known to work best when the reference scRNAseq dataset is of the same source as the bulk sample (Dong et al., 2020). Therefore we used a reference dataset which used the classic isolation method prior to scRNAseq (Strunz *et al*., 2020) to computationally deconvolve our bulk RNAseq data of the cell pellets from both isolation methods using bisqueRNA (Jew et al., 2020) (Figure 4A). To create a scRNAseq reference set which best matched our experimental setup, we only used untreated cells (i.e. healthy, PBS treated) from the reference dataset and kept the cellular level annotation described in the original reference dataset (Figure 4B). Next, we performed bisqueRNA deconvolution to explore differences in cell type predictions between the two isolation methods (Figure 4C). We found that distal progenitors isolated with the classic method were predicted to contain a small number of basal cells (see Table S2 for genes used to deconvolve each cell type) in comparison to no predicted basal cells in the 3DLD isolation (Figure 4D). Moreover, a higher proportion of other cell types such as neuroendocrine cells, known to be mainly located in the trachea and upper airways of mice, was also observed in cells isolated with the classic method (Figure 4D). This data supports the hypothesis that the classical isolation methods which rely on manual removal of the trachea and extrapulmonary airways may introduce unintentional contamination of proximal cells.

**Figure 4.**
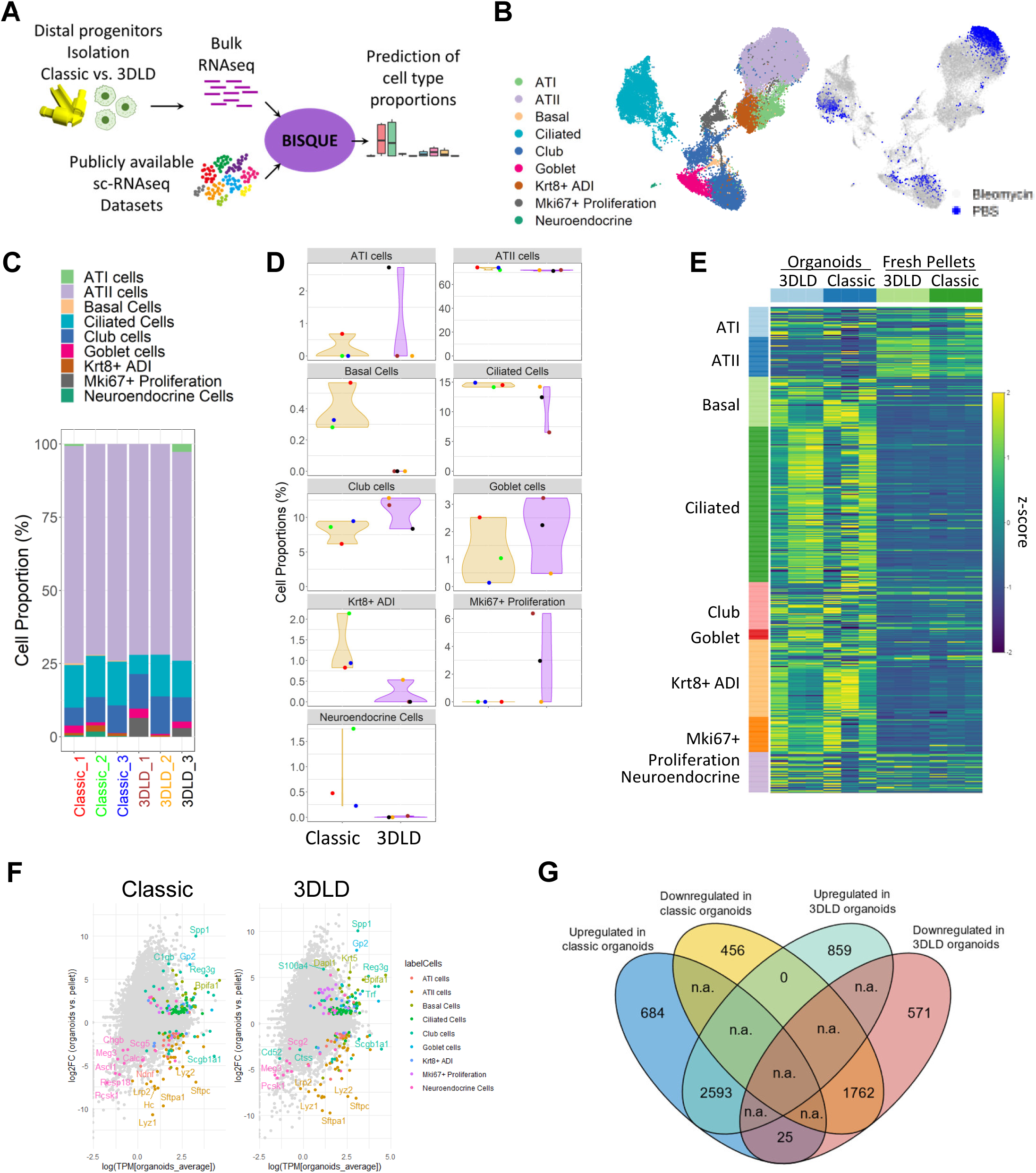
Deconvolution of bulk RNAseq of distal progenitor pellets predicts higher proportions of contaminating proximal airway cells in the classic isolation. (A) Approach for computational deconvolution with BisqueRNA. (B) UMAP of single cell dataset re-analyzed based on the original publication (Strunz *et al*., 2020). Only cells obtained from control mice (treated with PBS) were used in the deconvolution (marked in blue on the right UMAP). (C,D) Predicted epithelial cell proportion by cell type in the classic or 3DLD isolation method (C) and by individual isolation per mouse (D). (E) Heatmap of z-score of distal progenitors isolated with classic or 3DLD methods before and after differentiation in organoid culture (z-score: Gene(sample) – Gene(mean of all samples) / standard deviation). Genes are selected based on the markers of each cluster in the single cell dataset (Table S2). (F) Differential expression between organoids and pellets for both isolations colored by cell type markers (Table S2); only significant marker genes are highlighted (p-value <0.05; logFC>1) (Classic isolation: 5520 genes differentially expressed, of which 179 are marker genes) (3DLD isolation: 5810 genes differentially expressed, of which 211 are marker genes). (G) Venn diagram of overlapping upregulated and downregulated genes from differential expression analysis of organoids vs. pellets. Related to Figure S5 – S6.

Next, we explored transcriptional changes occurring within the organoids derived from both isolation methods. We first evaluated changes in phenotypic markers at the bulk RNAseq level across the different experimental conditions using z-scoring of gene lists which define each of the cell clusters from the reference single cell dataset (Table S2). Interestingly, noticed a shift towards increased expression of proximal epithelial markers, regardless of cell isolation method (Figure 4E) but with larger and significant increases in several basal cell and transitional cell (e.g. Krt8+) markers in cells isolated with the classic isolation method (Figure 4F). Next, we compared the transcriptional changes between the organoids and initial cell populations from which they were formed and found that while there were many genes in common which changed during organoid formation, there were 25 genes that were oppositely and significantly regulated (i.e. downregulated in 3DLD relative to 3DLD pellet and upregulated in classic organoids relative to classic pellets) (Figure 4G). In further exploring these genes, we found that they are expressed at various baseline levels and, interestingly, many are known to be involved in innate immunity, among other biological processes (Figure S5). Further, most of these genes were predominantly found in cells annotated as airway cells in the scRNAseq reference dataset, such as basal, ciliated, club, and neuroendocrine cells (Figure S6). As these organoids are cultured in the exact same *in vitro* conditions, our data indicates that the presence of rare cell populations due to the cell isolation method can have a significant impact on the transcriptional landscape within cultured organoids.

## Discussion

The lung epithelium is increasingly recognized as a central player in lung homeostasis and disease and a variety of techniques are employed for studying the potential of different epithelial cell progenitor populations in lung repair and regeneration. While lineage-tracing using reporter animals and scRNAseq approaches have been invaluable in describing this cellular diversity, the study of isolated and specific lung epithelial cell populations *in vitro* has been hampered by the lack of surface markers that distinguish the different cells along the respiratory epithelium. Additionally, reporter mice have been previously acknowledged to be ‘leaky’ and to label multiple cell types (Perl *et al*., 2009). This transcriptional heterogeneity has been confirmed by recent scRNAseq studies showing that some of the classically described phenotypic markers are expressed in multiple cell types along the entire airway epithelium (Strunz *et al*., 2020). Additionally, and especially in the case of disease, cells may both lose and/or acquire the phenotypic markers proposed to be used for their isolation. Therefore, the use of reporter mice and even surface markers is not sufficient alone for isolation of well-defined subpopulations along the airway epithelium.

In order to overcome these limitations, the majority of previous approaches for isolating either proximal or distal lung epithelial cells have used physical ligation between the proximal and distal epithelium at the level of the trachea or bronchi and isolated cells through dissociation via enzymatic digestion and selection using a) *in vitro* culture conditions in the case of proximal epithelial cells to enrich for basal progenitor cells or b) a combination of negative and positive surface markers for sorting of distal progenitor cells (Corti et al., 1996; Dobbs *et al*., 1986; Jansing *et al*., 2018; Lam *et al*., 2011; Vanderbilt *et al*., 2015; You *et al*., 2002). However, both techniques rely on precise surgical techniques to separate the proximal and distal epithelial compartments. Reproducible ligation of the trachea and bronchi is surgically challenging in mice and imprecise ligation or variability between laboratories in where and how ligation occurs may therefore inadvertently lead to the co-isolation of undesired cell populations at low frequency. This variability is particularly problematic considering current state-of-the-art techniques for studying lung repair and regeneration, such as organoid assays or scRNA-seq, which seek to identify and characterize the identity and behavior of rare cell populations *ex vivo*. Therefore, and to overcome the above-mentioned limitations, we designed and validated the use of a 3D printed guide to reproducibly and reliably isolate both proximal and distal progenitor cells from individual mice for use in multiple state-of-the-art *in vitro* assays such as ALI and organoid culture.

We found that this device allows for easy, quick, and reproducible ligation at the level of the left and right main bronchi, which is otherwise challenging and time consuming in mice. Further, we show that the minor modifications for cell isolation protocols introduced with the 3DLD for dual proximal and distal epithelial cell isolation does not impair the usage of isolated proximal or distal progenitor cells in downstream assays and analysis techniques. However, we found that distal epithelial cells freshly isolated with the 3DLD contained lower transcriptional variability as compared to those isolated with the classic method as well as lower amounts of proximal airway cell populations, as predicted by computational deconvolution. Taken together, our data indicates that the 3DLD helps improve reproducibility and the precision at which cell isolation occurs.

Heterogeneity in organoid size has been previously observed and while incompletely understood, has been shown to be altered due to changes in the initial epithelial cell population seeded into Matrigel as well as co-culture with different stromal cell populations or the presence of certain secreted factors in the media (Jain *et al*., 2015; Shiraishi et al., 2019). In this study, we did not perform additional selection of specific distal lung epithelial progenitor populations and thus our organoid culture theoretically contains multiple progenitor populations (Louie *et al*., 2022). Further studies are needed to understand whether the changes we observe in organoid size between the two isolation methods are due to secretion of paracrine factors altering distal progenitor cell organoid formation or directly from organoid formation from differences in the initial epithelial cell population.

Nonetheless, we found that most of the transcriptional changes which occurred at the bulk RNA-seq level during organoid formation in Matrigel were similar between the two isolation methods and that many of these changes were associated with the acquisition of markers associated with fully differentiated and mature lung epithelial cell types, including a diverse amount of phenotypic markers characteristic of proximal airway markers. We observed increases in classically described proximal airway markers, even in the 3DLD isolated distal epithelial cells where we observed little to no expression of these markers at baseline. The relevance of these changes and whether they can occur to this extent *in vivo* remains to be determined. It has recently been shown that *ex vivo* culture in Matrigel alters the transcriptional profile of the isolated progenitors at a single cell level and some of these changes are also observed when the progenitors are transplanted *in vivo* (Louie *et al*., 2022). However, this points to the increasingly recognized plasticity of epithelial cells observed *in vivo* and *ex vivo* in assays such as organoids, at least partially governed by the extracellular niche in which the cells reside in, comprised of both matrix and paracrine signaling (Loebel et al., 2021).

While organoids grown from the 3DLD or classic isolated cells changed similarly in relation to their pellets, a subset of genes which were significantly and differentially regulated in formed organoids. In particular, several genes that are associated with innate immunity were significantly upregulated in organoids formed from the classic method. These changes were reproducible in all organoids generated from individual mice in the classic isolation method, indicating that even though the organoids from both isolation methods were cultured in otherwise identical conditions, the composition of the initial cell population significantly impacted the trajectory and transcriptional landscape of the final organoid culture. The observation that organoids from classically isolated cells expressed higher transcriptional levels of several markers for innate immunity is particularly interesting as an increasing number of studies have shown a link between innate immunity and lung regeneration. Therefore, careful characterization of starting cell populations and the isolation methods used to obtain them are critical for interpreting the regenerative capacity of distal and proximal airway epithelial cells and the cues which govern their proliferation and differentiation.

In conclusion, we present a method to simultaneously isolate murine proximal and distal epithelial progenitors from the same mouse lung in a reproducible and inexpensive manner utilizing a 3D printed guide. This method allows the evaluation of both the distal and proximal epithelial progenitor populations from a single animal and will allow for increased reproducibility across laboratories performing state-of-the-art *ex vivo* analysis for regenerative populations in diverse downstream analyses such as scRNAseq, organoid assays, or other 3D *in vitro* assays.

## Experimental procedures

All media formulations are listed in table S1.

### Animals

The use of mice was approved by the Local Ethics Committee for Animal Research in Lund, Sweden (Dnr 6436/2017). All animal experiments were conducted in accordance with legislation by the European Union (2010/63/EU) and local Swedish law. All animals received care according to the Guide for the Care and Use of Laboratory Animals: Eighth Edition, National Academies Press (2011). 8 – 12 weeks C57BL/6N male mice (*Mus Musculus*) were used for all isolations of this study. Mice were housed with same-sex litter mates under alternating light-dark cycles in a humidity- and temperature-controlled room with food and water *ad libitum*. All mice were provided care in accordance with the animal protection and welfare guidelines, and experiments and data are reported in accordance with ARRIVE 2.0 guidelines.

### Fused Deposition Modelling (FDM) 3D printing

3D lobe divider (3DLD) was designed using Blender Software (V2.8, www.blender.org). The 3D model is available on GitHub Repository (https://github.com/Lung-bioengineering-regeneration-lab/dual_cell_isolation/)(Figure 1B).

### Murine lung explantation and preparation for 3DLD or classic distal isolation methods

All mice were euthanized using a 3:1 ketamine/xylazine mixture and their tracheas were intubated using a modified blunt 19-gauge needle. The lungs were then perfused through the heart with PBS, warmed to 37 °C, and removed by dissecting the esophagus and connective tissue away. Lungs were immersed in PBS at room temperature (RT) to remove any residual blood. For the 3DLD-based method, lungs were placed in the 3D printed guide by placing the trachea within the main channel and stabilized with sutures. Loose sutures were placed around both the right and left bronchi in preparation for instillation of dissociation solutions and to allow for rapid ligation (Figure 1C).

### 3DLD and classic distal cell isolation and organoid formation

For both distal cell isolation procedures, distal dissociation solution (Table S1) was injected through the cannulated trachea. In 3DLD isolations, this was followed by immediate closure of the 3DLD device and knotting the sutures surrounding the main bronchi. The lobes were then separated proximally to the surgical knot with scissors and placed on ice until proceeding to cell isolation. The trachea was removed and immediately washed with cold Ham’s F12/AB Medium and placed in Ham’s F12/AB for further processing (Table S1). For the classic isolation, ligation occurred manually on the trachea after injection of the distal dissociation solution. The lungs were placed on ice until proceeding to cell isolation. Distal progenitor cells for both the classic and 3DLD device were isolated as previously described (Jansing *et al*., 2018). Distal organoid formation was performed with CCL206 cells (ATCC), murine lung fibroblasts (Mlg), as stromal support cells previously described (Costa et al., 2021). Further details for cell isolation and organoid formation, collection, fixation and imaging can be found in supplemental experimental procedures.

### Proximal progenitor isolation and expansion

Proximal progenitors were isolated as previously described with minor modifications (Eenjes et al., 2018; Lam *et al*., 2011). Connective tissue was removed from 3DLD isolated tracheas and the inner lumen was exposed by cutting the trachea open with surgical scissors. Opened tracheas from individual mice were incubated in F12/pronase overnight at 4 °C (Table S1) then washed in F12/AB 3 times to collect digested cells. Cells from each individual mouse were pelleted and treated with 500 μg/ml DNase I (A3778, Saveen & Werner, Sweden) for 5 minutes on ice. Cells isolated from individual mice were then expanded separately in proximal progenitor expansion medium (Table S1). When cells reached 80-90 % confluence, they were passaged and either seeded submerged, in air liquid interface (ALI) culture, or in Matrigel for organoid formation as described in supplemental experimental procedures.

### Total RNA isolation and Next Generation Sequencing

RNA isolation was performed using the RNeasy Micro Kit (74004, QIAGEN, Sweden) including RNase-free DNase I incubation (79254, QIAGEN) following the manufacturer’s protocol. RNA concentrations were measured using NanoDrop 1000 and Qubit™ RNA HS Assay Kit (Q32852, Fisher Scientific). RNA quality was evaluated by the Agilent 2100 BioAnalyzer and sequenced using NovaSeq 6000 Sequencing System (20012850, Illumina) as described in supplemental experimental procedures.

### Deconvolution of bulk RNAseq data using publicly available datasets

Bulk RNAseq data was computationally deconvolved using BisqueRNA to predict the proportions of epithelial cells in our two isolation methods (Jew *et al*., 2020). The reference single cell dataset is comprised of lung and airway epithelial cells isolated using the classic isolation method for distal lung cells and is available at the Gene Expression Omnibus (accession code GSE141259) (Strunz *et al*., 2020). Bulk RNAseq read counts from distal cells of the 3DLD isolation and classic isolation were filtered based on gene expression; only genes with counts in at least 2 out of the 3 samples in each of the groups were considered. ExpressionSets were generated from the Bulk Data and single cell data based on the original single cell reference dataset annotation using Seurat (Butler et al., 2018; Strunz *et al*., 2020). BisqueRNA deconvolution was done using the function *ReferenceBasedDecomposition* with options of *use*.*overlap* and *old*.*cpm* selected as False. All code used is available on the GitHub Repository (https://github.com/Lung-bioengineering-regeneration-lab/dual_cell_isolation)

### Statistical Analysis

Statical tests were done using R statistical software and applied where appropriate and are indicated in figure legends. P-value cut-offs are indicated in each figure.

### Resource availability

Raw data from the current study is deposited in the European Nucleotide Archive (ENA) under accession no. E-MTAB-11502. All relevant codes and 3D designs used in this study are available in the GitHub Repository (https://github.com/Lung-bioengineering-regeneration-lab/dual_cell_isolation).

## Acknowledgments

We thank the Lund University Bioimaging Centre (LBIC) for their assistance in various imaging techniques. We thank visiting Ph.D. student Caroline Pereira for insightful conversations and assistance with cell isolation. We thank the Center for Translational Genomics, Lund University and Clinical Genomics Lund, SciLifeLab for providing sequencing services. This project is partially funded by the Swedish Research Council Starting Grant (Dnr 2018-02352). The Knut and Alice Wallenberg foundation is acknowledged for generous support. This project has received funding from the European Research Council (ERC) under the European Union’s Horizon 2020 research and innovation programme (grant agreement No 805361). The LSFM imaging was enabled by resources funded in part by the Crafoord Foundation (Dnr 20200744). Computation/data handling was enabled by resources provided by the Swedish National Infrastructure for Computing (SNIC) at LUNARC partially funded by the Swedish Research Council through grant agreement no. 2018-05973.

## Author Contributions

Conceptualization: HNA, VP, LM, DEW; Data curation: HNA; Formal analysis: HNA, DEW; Funding acquisition: DEW, JS, LM; Investigation: HNA, JS, IAS, MM; Methodology: HNA, JS, IAS, DEW; Project administration: HNA, VP, DEW; Resources and Software: HNA, DEW; Supervision: JS, DEW; Validation and visualization: HNA; Writing original manuscript: HNA, DEW; Review and editing: All authors.

## References

Barkauskas, C.E., Chung, M.-I., Fioret, B., Gao, X., Katsura, H., and Hogan, B.L.M. (2017). Lung organoids: current uses and future promise. Development 144, 986–997. 10.1242/dev.140103.

Barkauskas, C.E., Cronce, M.J., Rackley, C.R., Bowie, E.J., Keene, D.R., Stripp, B.R., Randell, S.H., Noble, P.W., and Hogan, B.L.M. (2013). Type 2 alveolar cells are stem cells in adult lung. The Journal of Clinical Investigation 123, 3025–3036. 10.1172/JCI68782.

Butler, A., Hoffman, P., Smibert, P., Papalexi, E., and Satija, R. (2018). Integrating single-cell transcriptomic data across different conditions, technologies, and species. Nature Biotechnology 36, 411–420. 10.1038/nbt.4096.

Chapman, H.A., Li, X., Alexander, J.P., Brumwell, A., Lorizio, W., Tan, K., Sonnenberg, A., Wei, Y., and Vu, T.H. (2011). Integrin α6β4 identifies an adult distal lung epithelial population with regenerative potential in mice. J Clin Invest 121, 2855–2862. 10.1172/jci57673.

Chen, H., Matsumoto, K., Brockway, B.L., Rackley, C.R., Liang, J., Lee, J.-H., Jiang, D., Noble, P.W., Randell, S.H., Kim, C.F., and Stripp, B.R. (2012). Airway epithelial progenitors are region specific and show differential responses to bleomycin-induced lung injury. Stem cells (Dayton, Ohio) 30, 1948–1960. 10.1002/stem.1150.

Corti, M., Brody, A.R., and Harrison, J.H. (1996). Isolation and primary culture of murine alveolar type II cells. American journal of respiratory cell and molecular biology 14, 309–315. 10.1165/ajrcmb.14.4.8600933.

Costa, R., Wagner, D.E., Doryab, A., De Santis, M.M., Schorpp, K., Rothenaigner, I., Lehmann, M., Baarsma, H.A., Liu, X., Schmid, O., et al. (2021). A drug screen with approved compounds identifies amlexanox as a novel Wnt/β-catenin activator inducing lung epithelial organoid formation. British Journal of Pharmacology 178, 4026–4041. https://doi.org/10.1111/bph.15581.

Danto, S.I., Shannon, J.M., Borok, Z., Zabski, S.M., and Crandall, E.D. (1995). Reversible transdifferentiation of alveolar epithelial cells. American journal of respiratory cell and molecular biology 12, 497–502. 10.1165/ajrcmb.12.5.7742013.

Dobbs, L.G., Gonzalez, R., and Williams, M.C. (1986). An Improved Method for Isolating Type II Cells in High Yield and Purity. 134, 141–145. 10.1164/arrd.1986.134.1.141.

Dong, M., Thennavan, A., Urrutia, E., Li, Y., Perou, C.M., Zou, F., and Jiang, Y. (2020). SCDC: bulk gene expression deconvolution by multiple single-cell RNA sequencing references. Briefings in Bioinformatics 22, 416–427. 10.1093/bib/bbz166.

Eenjes, E., Mertens, T.C.J., Buscop-van Kempen, M.J., van Wijck, Y., Taube, C., Rottier, R.J., and Hiemstra, P.S. (2018). A novel method for expansion and differentiation of mouse tracheal epithelial cells in culture. Scientific Reports 8, 7349. 10.1038/s41598-018-25799-6.

Jain, R., Barkauskas, C.E., Takeda, N., Bowie, E.J., Aghajanian, H., Wang, Q., Padmanabhan, A., Manderfield, L.J., Gupta, M., Li, D., et al. (2015). Plasticity of Hopx(+) type I alveolar cells to regenerate type II cells in the lung. Nat Commun 6, 6727. 10.1038/ncomms7727.

Jansing, N.L., McClendon, J., Kage, H., Sunohara, M., Alvarez, J.R., Borok, Z., and Zemans, R.L. (2018). Isolation of Rat and Mouse Alveolar Type II Epithelial Cells. In Lung Innate Immunity and Inflammation: Methods and Protocols, S. Alper, and W.J. Janssen, eds. (Springer New York), pp. 69–82. 10.1007/978-1-4939-8570-8_6.

Jew, B., Alvarez, M., Rahmani, E., Miao, Z., Ko, A., Garske, K.M., Sul, J.H., Pietiläinen, K.H., Pajukanta, P., and Halperin, E. (2020). Accurate estimation of cell composition in bulk expression through robust integration of single-cell information. Nature Communications 11, 1971. 10.1038/s41467-020-15816-6.

Joshi, N., Watanabe, S., Verma, R., Jablonski, R.P., Chen, C.-I., Cheresh, P., Markov, N.S., Reyfman, P.A., McQuattie-Pimentel, A.C., Sichizya, L., et al. (2019). A spatially restricted fibrotic niche in pulmonary fibrosis is sustained by M-CSF/M-CSFR signaling in monocyte-derived alveolar macrophages. European Respiratory Journal, 1900646. 10.1183/13993003.00646-2019.

Kim, C.F.B., Jackson, E.L., Woolfenden, A.E., Lawrence, S., Babar, I., Vogel, S., Crowley, D., Bronson, R.T., and Jacks, T. (2005). Identification of Bronchioalveolar Stem Cells in Normal Lung and Lung Cancer. Cell 121, 823–835. https://doi.org/10.1016/j.cell.2005.03.032.

Lam, H.C., Choi, A.M.K., and Ryter, S.W. (2011). Isolation of mouse respiratory epithelial cells and exposure to experimental cigarette smoke at air liquid interface. Journal of visualized experiments : JoVE, 2513. 10.3791/2513.

Loebel, C., Weiner, A.I., Katzen, J.B., Morley, M.P., Bala, V., Cardenas-Diaz, F.L., Davidson, M.D., Shiraishi, K., Basil, M.C., Ochs, M., et al. (2021). Microstructured hydrogels to guide self-assembly and function of lung alveolospheres. bioRxiv, 2021.2008.2030.457534. 10.1101/2021.08.30.457534.

Louie, S.M., Moye, A.L., Wong, I.G., Lu, E., Shehaj, A., Garcia-de-Alba, C., Ararat, E., Raby, B.A., Lu, B., Paschini, M., et al. (2022). Progenitor potential of lung epithelial organoid cells in a transplantation model. Cell Reports 39, 110662. https://doi.org/10.1016/j.celrep.2022.110662.

McQualter, J.L., Yuen, K., Williams, B., and Bertoncello, I. (2010). Evidence of an epithelial stem/progenitor cell hierarchy in the adult mouse lung. Proc Natl Acad Sci U S A 107, 1414–1419. 10.1073/pnas.0909207107.

Messier, E.M., Mason, R.J., and Kosmider, B. (2012). Efficient and rapid isolation and purification of mouse alveolar type II epithelial cells. Experimental Lung Research 38, 363–373. 10.3109/01902148.2012.713077.

Montoro, D.T., Haber, A.L., Biton, M., Vinarsky, V., Lin, B., Birket, S.E., Yuan, F., Chen, S., Leung, H.M., Villoria, J., et al. (2018). A revised airway epithelial hierarchy includes CFTR-expressing ionocytes. Nature 560, 319–324. 10.1038/s41586-018-0393-7.

Perl, A.-K., Zhang, L., and Whitsett, J.A. (2009). Conditional Expression of Genes in the Respiratory Epithelium in Transgenic Mice. American journal of respiratory cell and molecular biology 40, 1–3. 10.1165/rcmb.2008-0011ED.

Rawlins, E.L. (2008). Lung Epithelial Progenitor Cells. Proceedings of the American Thoracic Society 5, 675–681. 10.1513/pats.200801-006AW.

Rawlins, E.L., Okubo, T., Xue, Y., Brass, D.M., Auten, R.L., Hasegawa, H., Wang, F., and Hogan, B.L. (2009). The role of Scgb1a1+ Clara cells in the long-term maintenance and repair of lung airway, but not alveolar, epithelium. Cell Stem Cell 4, 525–534. 10.1016/j.stem.2009.04.002.

Reynolds, S.D., Giangreco, A., Power, J.H.T., and Stripp, B.R. (2000). Neuroepithelial Bodies of Pulmonary Airways Serve as a Reservoir of Progenitor Cells Capable of Epithelial Regeneration. The American Journal of Pathology 156, 269–278. https://doi.org/10.1016/S0002-9440(10)64727-X.

Riemondy, K.A., Jansing, N.L., Jiang, P., Redente, E.F., Gillen, A.E., Fu, R., Miller, A.J., Spence, J.R., Gerber, A.N., Hesselberth, J.R., and Zemans, R.L. (2019). Single cell RNA sequencing identifies TGFβ as a key regenerative cue following LPS-induced lung injury. JCI insight 5, e123637. 10.1172/jci.insight.123637.

Rock, J.R., Onaitis, M.W., Rawlins, E.L., Lu, Y., Clark, C.P., Xue, Y., Randell, S.H., and Hogan, B.L.M. (2009). Basal cells as stem cells of the mouse trachea and human airway epithelium. 106, 12771–12775. 10.1073/pnas.0906850106 %J Proceedings of the National Academy of Sciences.

Rock, J.R., Randell, S.H., and Hogan, B.L. (2010). Airway basal stem cells: a perspective on their roles in epithelial homeostasis and remodeling. Dis Model Mech 3, 545–556. 10.1242/dmm.006031.

Shiraishi, K., Shichino, S., Ueha, S., Nakajima, T., Hashimoto, S., Yamazaki, S., and Matsushima, K. (2019). Mesenchymal-Epithelial Interactome Analysis Reveals Essential Factors Required for Fibroblast-Free Alveolosphere Formation. iScience 11, 318–333. https://doi.org/10.1016/j.isci.2018.12.022.

Sikkema, L., Strobl, D., Zappia, L., Madissoon, E., Markov, N., Zaragosi, L., Ansari, M., Arguel, M., Apperloo, L., Bécavin, C., et al. (2022). An integrated cell atlas of the human lung in health and disease. bioRxiv, 2022.2003.2010.483747. 10.1101/2022.03.10.483747.

Strunz, M., Simon, L.M., Ansari, M., Kathiriya, J.J., Angelidis, I., Mayr, C.H., Tsidiridis, G., Lange, M., Mattner, L.F., Yee, M., et al. (2020). Alveolar regeneration through a Krt8+ transitional stem cell state that persists in human lung fibrosis. Nature Communications 11, 3559. 10.1038/s41467-020-17358-3.

Succony, L., Gómez-López, S., Pennycuick, A., Alhendi, A.S.N., Davies, D., Clarke, S.E., Gowers, K.H.C., Wright, N.A., Jensen, K.B., and Janes, S.M. (2021). <em>Lrig1</em> expression identifies airway basal cells with high proliferative capacity and restricts lung squamous cell carcinoma growth. European Respiratory Journal, 2000816. 10.1183/13993003.00816-2020.

Tata, P.R., and Rajagopal, J. (2017). Plasticity in the lung: making and breaking cell identity. Development 144, 755–766. 10.1242/dev.143784.

Vanderbilt, J.N., Gonzalez, R.F., Allen, L., Gillespie, A., Leaffer, D., Dean, W.B., Chapin, C., and Dobbs, L.G. (2015). High-efficiency type II cell-enhanced green fluorescent protein expression facilitates cellular identification, tracking, and isolation. American journal of respiratory cell and molecular biology 53, 14–21. 10.1165/rcmb.2014-0348MA.

Vaughan, A.E., Brumwell, A.N., Xi, Y., Gotts, J.E., Brownfield, D.G., Treutlein, B., Tan, K., Tan, V., Liu, F.C., Looney, M.R., et al. (2014). Lineage-negative progenitors mobilize to regenerate lung epithelium after major injury. Nature 517, 621. 10.1038/nature14112

https://www.nature.com/articles/nature14112#supplementary-information.

Vaughan, A.E., Brumwell, A.N., Xi, Y., Gotts, J.E., Brownfield, D.G., Treutlein, B., Tan, K., Tan, V., Liu, F.C., Looney, M.R., et al. (2015). Lineage-negative progenitors mobilize to regenerate lung epithelium after major injury. Nature 517, 621–625. 10.1038/nature14112.

Volckaert, T., and De Langhe, S. (2014). Lung epithelial stem cells and their niches: Fgf10 takes center stage. Fibrogenesis & Tissue Repair 7, 8. 10.1186/1755-1536-7-8.

You, Y., Richer, E.J., Huang, T., and Brody, S.L. (2002). Growth and differentiation of mouse tracheal epithelial cells: selection of a proliferative population. American journal of physiology. Lung cellular and molecular physiology 283, L1315–1321. 10.1152/ajplung.00169.2002.

Zuo, W., Zhang, T., Wu, D.Z.A., Guan, S.P., Liew, A.-A., Yamamoto, Y., Wang, X., Lim, S.J., Vincent, M., Lessard, M., et al. (2015). p63+Krt5+ distal airway stem cells are essential for lung regeneration. Nature 517, 616–620. 10.1038/nature13903.

