## Supplemental for "Simultaneous isolation of proximal and distal lung progenitor cells from individual mice using a 3D printed guide reduces proximal cell contamination of distal lung epithelial cell isolations"

**Figure S1. Differentiation of 3DLD isolated proximal epithelial cells into a pseudostratified epithelium with normal apical-basal polarity.** (A) Hematoxylin and eosin staining and bright field imaging of differentiated proximal epithelial cells cultured in ALI at day 28 confirming the presence of a pseudostratified monolayer. Imaged at 40X magnification (B) Immunofluorescence staining of cytokeratin5 (Krt5, apical layer), alpha-tubulin (α-Tub, cilia in the apical layer). Related to Figure 2. Scalebar: 50 μm.

**A**

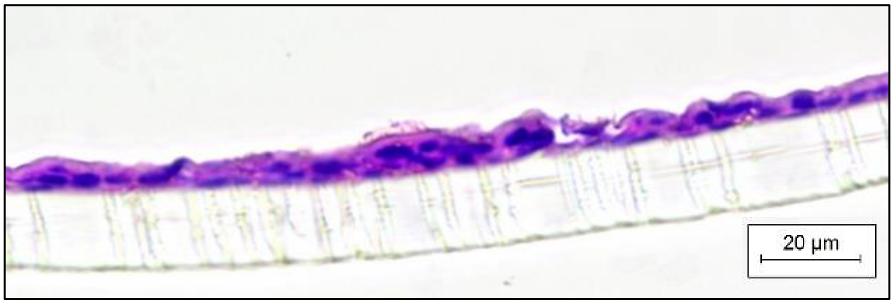


**B**

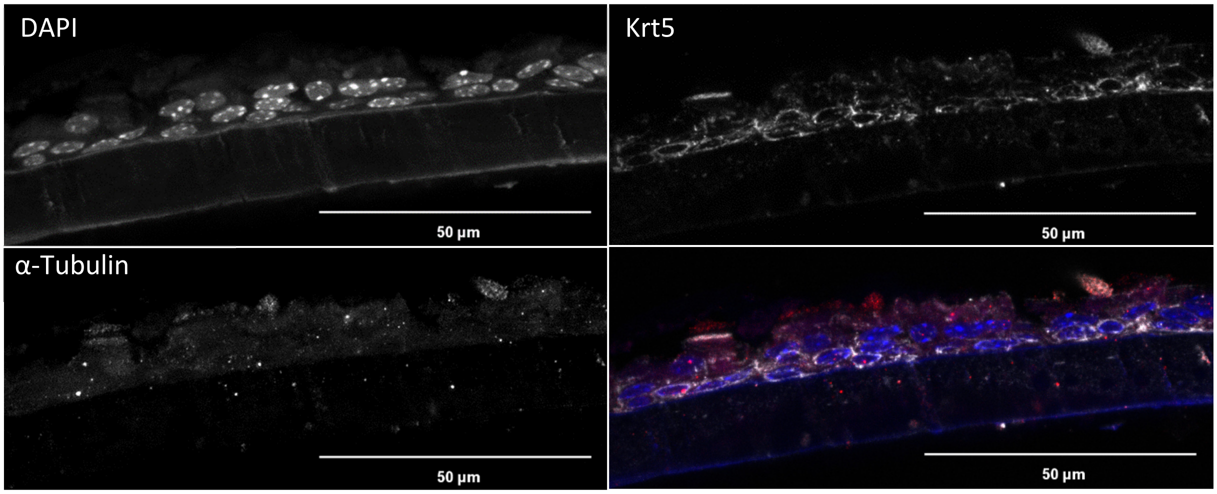

**Figure S2. Scanning electron microscopy of differentiated mouse proximal progenitor cells after 28 days of air liquid interface culture. Related to Figure 2.**

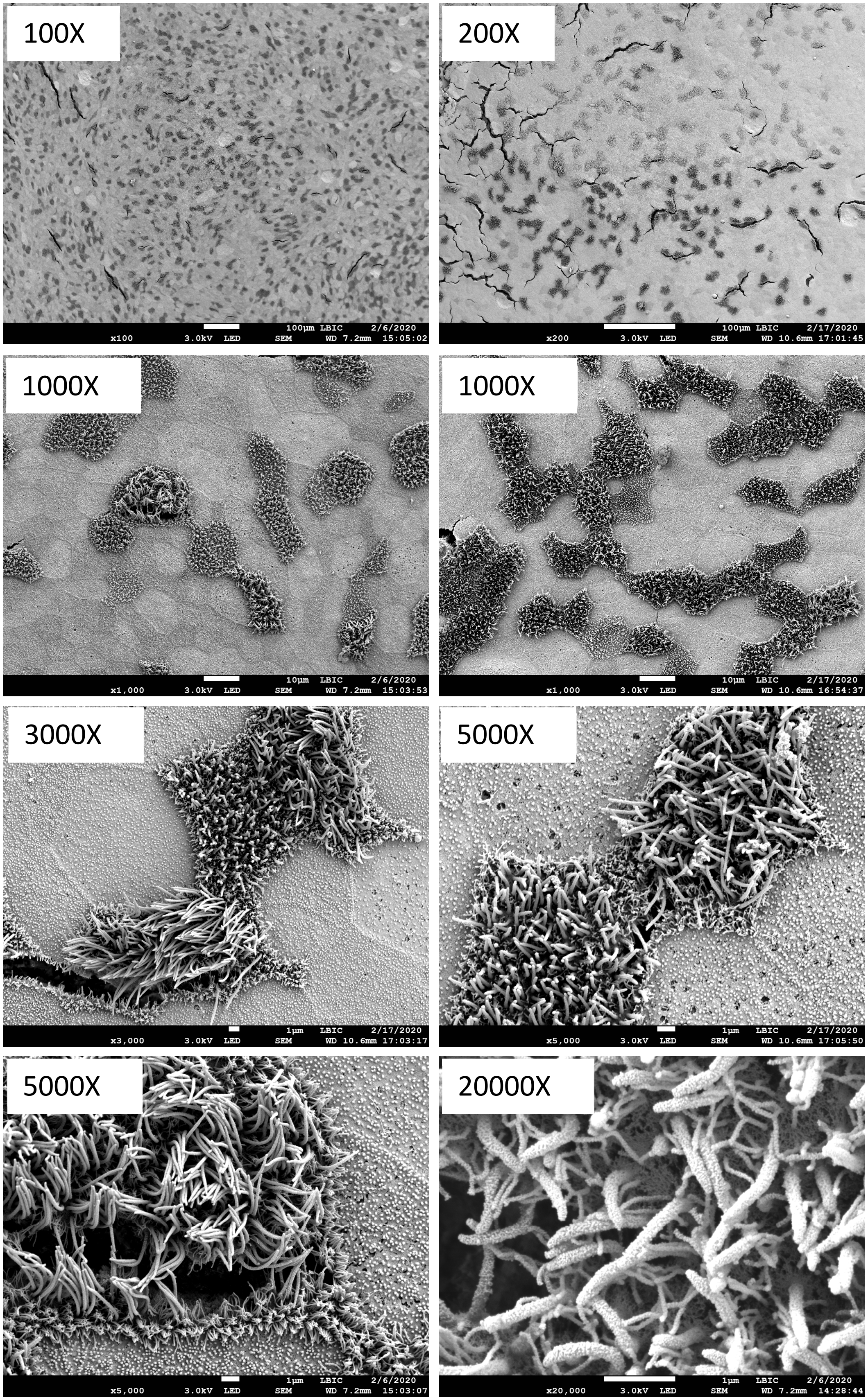

**Figure S3. Lightsheet fluorescent microscopy imaging of organoids grown from distal progenitors isolated using either the classic or 3DLD methods.** (A) Representative 2D images of different regions of interest (ROI) obtained from the z-stacks to display the nuclear (DAPI) staining of different size organoids. (B) Maximum intensity projection of the DAPI staining of organoids followed by segmentation of each organoid using virtual reality (VR) software SyGlass. These images were obtained at 0.4μm z-resolution which would not be otherwise possible with another imaging modality without photobleaching the sample.

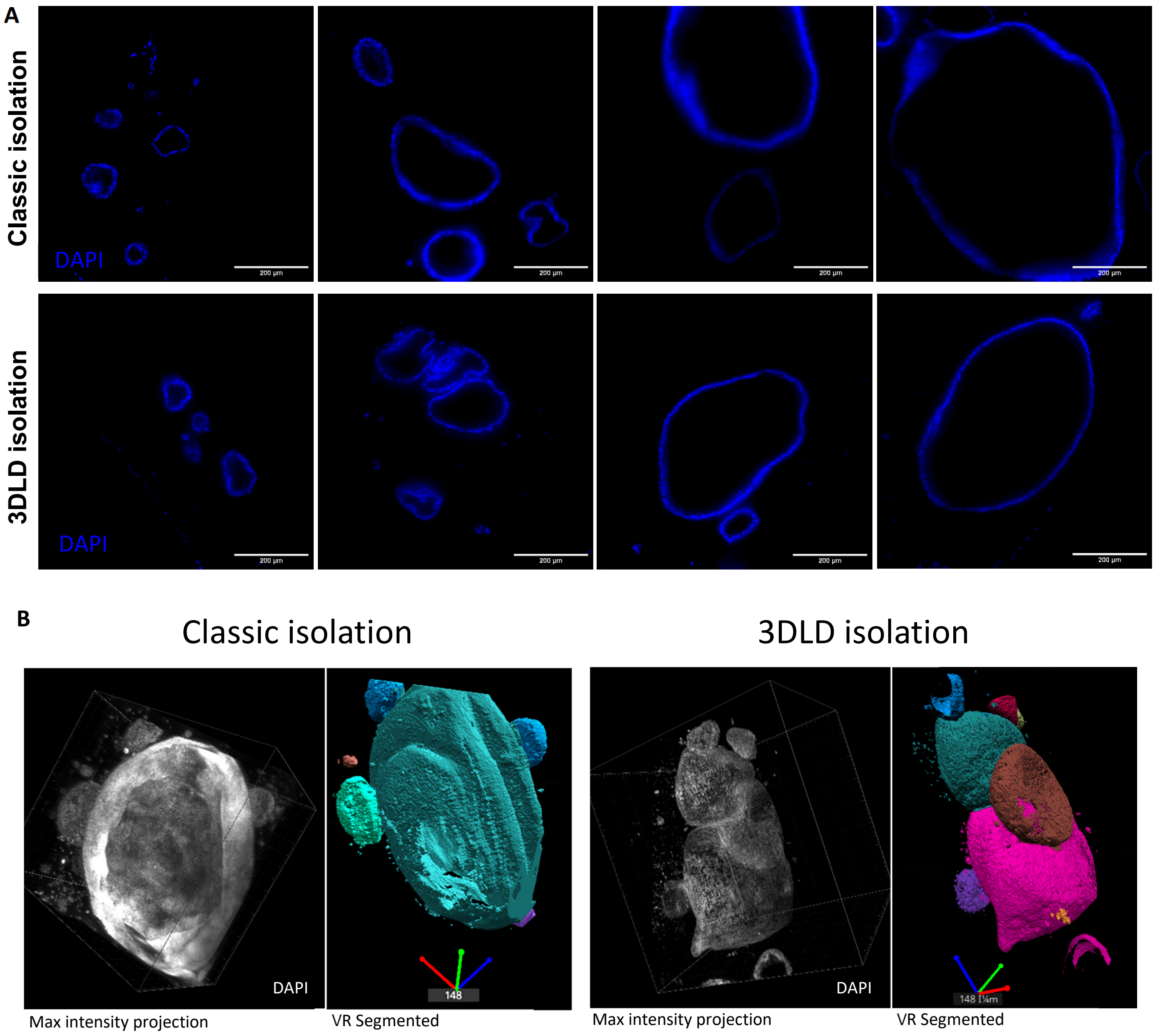

**Figure S4. Direct comparison of the relative variance for each gene**

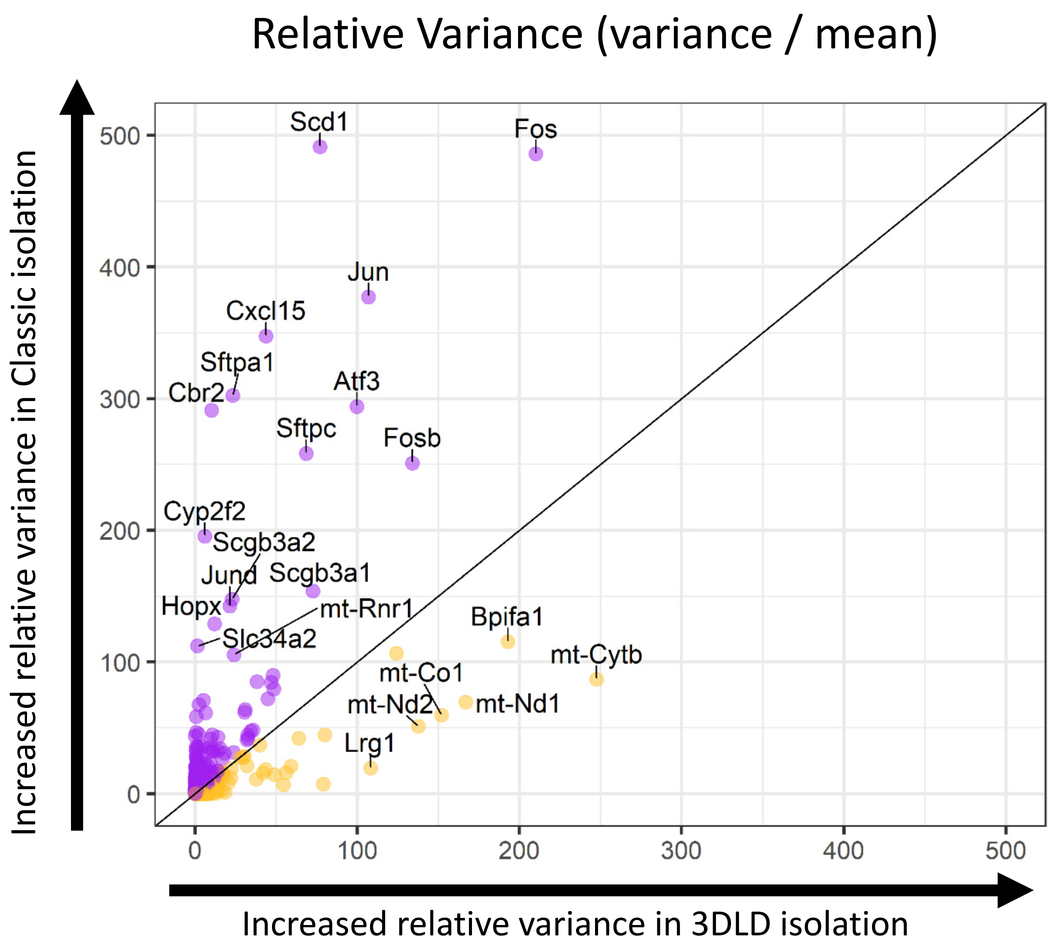

**Figure S5. Gene expression of genes that are oppositely and significantly regulated along the expression trajectory of organoid culture (i.e. downregulated in 3DLD relative to 3DLD pellet and upregulated in classic organoids relative to classic pellets). Related to Figure 4.** Data shown in transcripts per million (TPM).

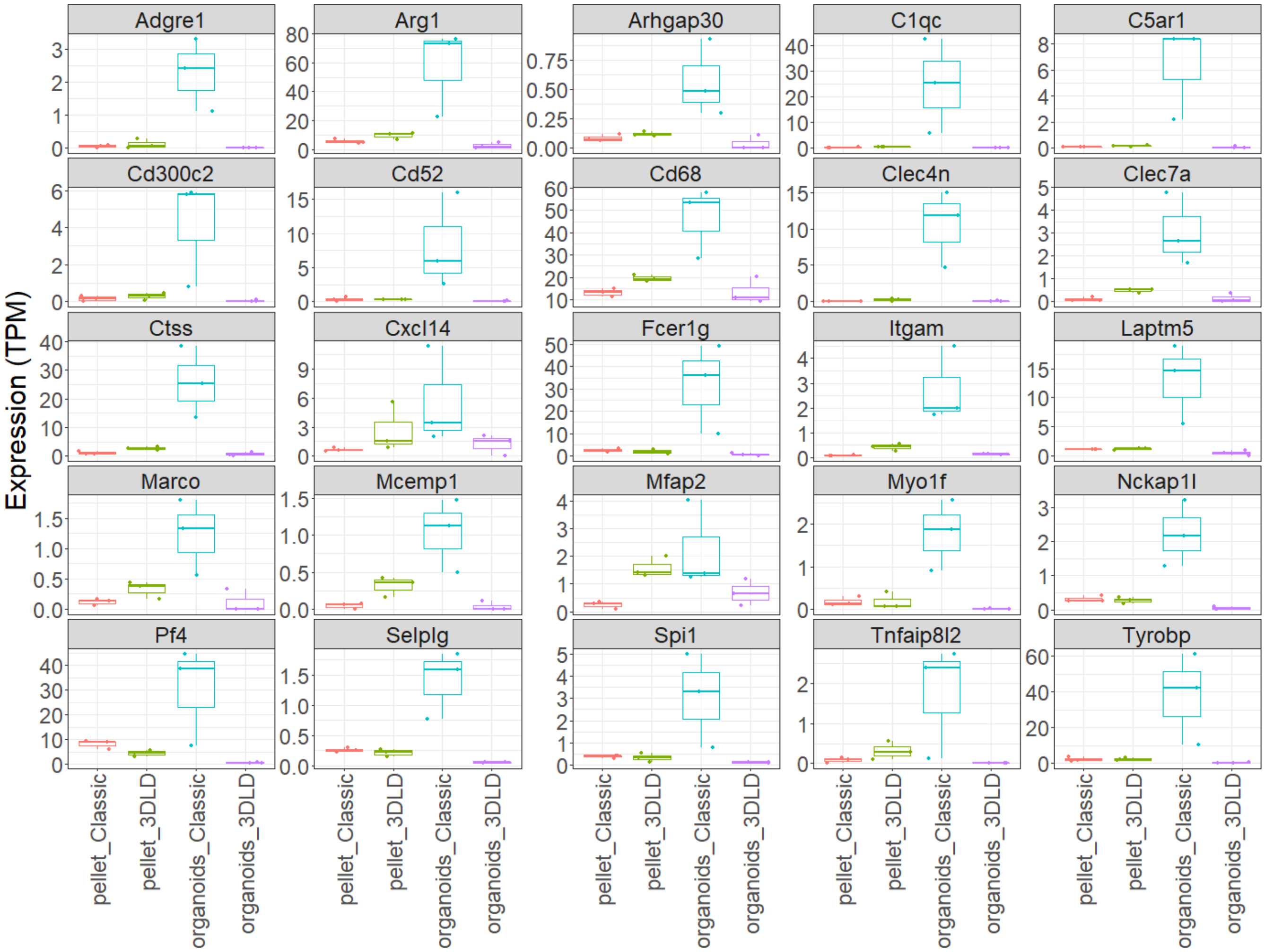

**Figure S6: FeaturePlots of genes indicated in Figure S4 projected on publicly available single cell Dataset (GSE141259) of epithelial cells isolated in a similar manner to the classic isolation. Related to Figure 4.** Scaled data is used to generate the featureplots using the Seurat package on R. Scale bar is relative for each feature.

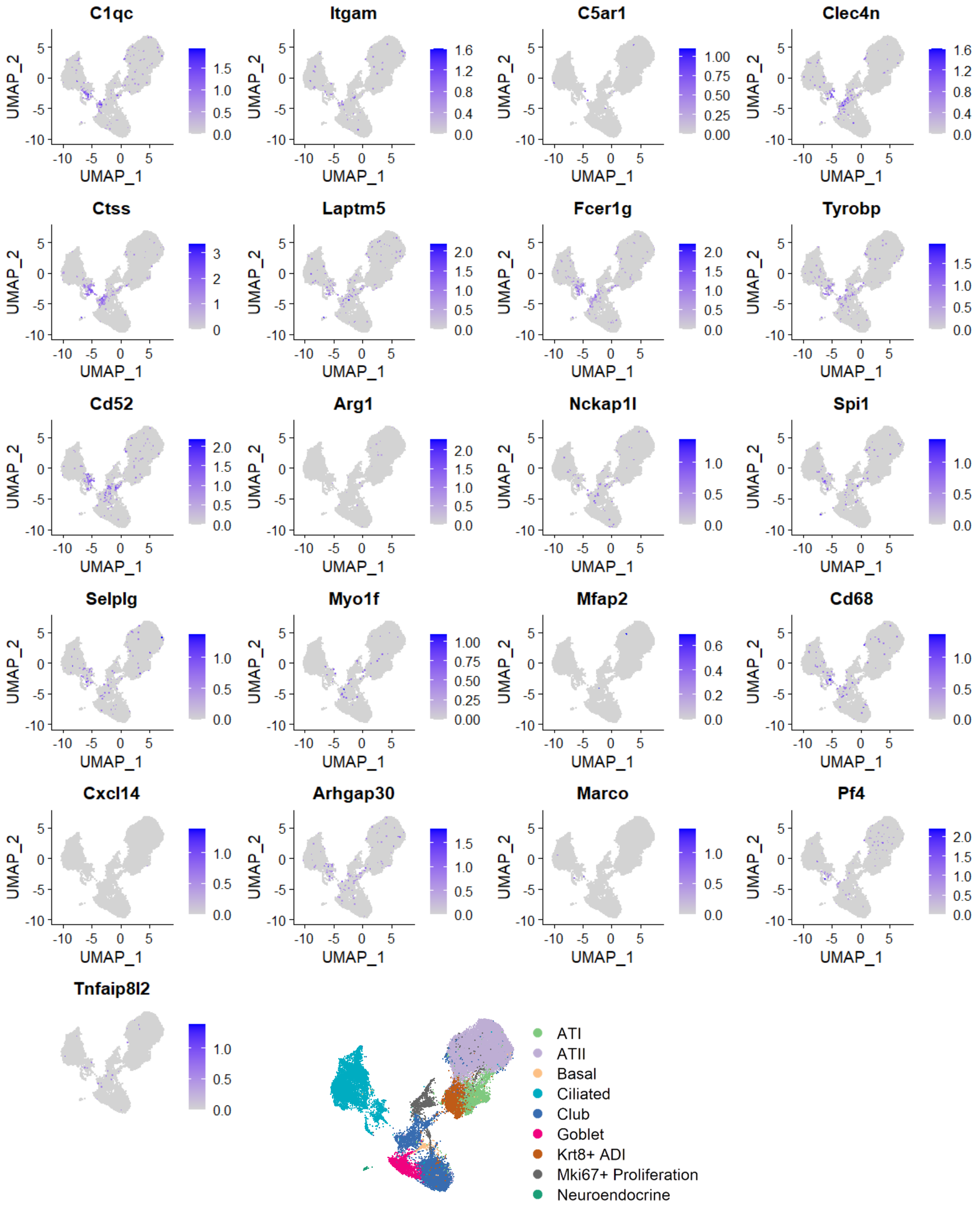

**Table S1: Composition of all buffers and solutions.**

| Cell type | Medium | Component | Distributor (product #) | Final Concentration |
|---|---|---|---|---|
| Distal progenitor s | Distal dissociation solution | Dispase | Fisher Scientific Sweden (11553550) | 50 caseinolytic units/ml |
| | Distal base medium | DMEM | Sigma-Aldrich (D5546) | - |
| | | D(+)-glucose Anhydrous | Saveen & Werner (141341.1211) | 3.6 mg/ml |
| | | GlutaMAX | Fisher Scientific Sweden (13462629) | 2 % |
| | | Pen/Strep | Fisher Scientific Sweden (15140122) | 100 U/ml, 100 µg/ml |
| | | HEPES | Fisher Scientific Sweden (11464805) | 10 mM |
| | Base/DNase | Distal Base Medium | *Described above* | - |
| | | DNASE I | Saveen & Werner (A3778)<br>Specific lot contained 3441.4 U/mg | 0.04 mg/ml |
| | Distal organoid medium | DMEM/F12 | Fisher Scientific Sweden (11554546) | - |
| | | GlutaMAX | Fisher Scientific Sweden (13462629) | 1X |
| | | Pen/Strep | Fisher Scientific Sweden (15140122) | 100 U/ml, 100 µg/ml |
| | | Amphotericin B | Fisher Scientific Sweden (11520496) | 1X |
| | | Insulin-Transferrin-Selenium (ITS) | Fisher Scientific Sweden (12097549) | 1X |
| | | Cholera Toxin | Sigma-Aldrich (C8052-.5MG) | 0.1 µg/ml |
| | | Murine EGF | PeproTech (315-09-100ug) | 25 ng/ml |
| | | BPE | Fisher Scientific Sweden (11568866) | 30 µg/ml |
| | | Retinoic Acid (Freshly added) | Sigma-Aldrich (R2625-50MG) | 0.01 µM |
| | | Y-27632 (only first 48 hours) | Cayman Chemical Company (10005583) | 10 µM |
| CCL206 | CCL206 culture medium | DMEM/F12 | Fisher Scientific Sweden (11554546) | - |
| | | Pen/Strep | Fisher Scientific Sweden (15140122) | 100 U/ml, 100 µg/ml |
| | | FBS | Fisher Scientific Sweden (11573397) | 10 % |
| Proximal progenitor s | Ham's F12/AB | Ham's F12 | Fisher Scientific Sweden (15172529) | - |
| | | Pen/Strep | Fisher Scientific Sweden (15140122) | 100 U/ml, 100 µg/ml |
| | | Amphotericin B | Fisher Scientific Sweden (11520496) | 1X |
| | F12/pronase | Ham's F12/AB | | |
| | | Pronase | Sigma Aldrich (P5147) | 1.5 mg/ml |
| | Proximal progenitor expansion medium | Kerratinocyte-SFM with L-gluatmine | Fisher Scientific Sweden (11580526) | - |
| | | Pen/Strep | Fisher Scientific Sweden (15140122) | 100 U/ml, 100 µg/ml |
| | | Murine EGF | PeproTech (315-09-100ug) | 0.025 µg/ml |
| | | Bovine Pituitary Extract (BPE) | Fisher Scientific Sweden (11568866) | 0.03 mg/ml |
| | | (-)-Isoproterenol hydrochloride | Sigma-Aldrich (I6504-100MG) | 1 µM |
| | | Y-27632 | Cayman Chemical Company (10005583) | 10 µM |
| | | DAPT | Sigma-Aldrich (D5942-5MG) | 5 µM |
| | Proximal B | DMEM/F12 | Fisher Scientific Sweden (11554546) | - |
| | | Pen/Strep | Fisher Scientific Sweden (15140122) | 100 U/ml, 100 µg/ml |
| | | Amphotericin B | Fisher Scientific Sweden (11520496) | 1X |
| | | NaHCO3 | Fisher Scientific Sweden (10308462) | 0.03 % |

| | | | | |
|---|---|---|---|---|
| | | HEPES | Fisher Scientific Sweden (11464805) | 15 mM |
| | Proximal progenitor proliferation medium | Proximal B | *Described above* | - |
| | | L-Glutamine | Fisher Scientific Sweden (12569079) | 1.5 mM |
| | | ITS | Fisher Scientific Sweden (12097549) | 1X |
| | | Cholera Toxin | Sigma-Aldrich (C8052-.5MG) | 0.1 µg/ml |
| | | Murine EGF | PeproTech (315-09-100ug) | 0.025 µg/ml |
| | | BPE | Fisher Scientific Sweden (11568866) | 0.03 mg/ml |
| | | Y-27632 | Cayman Chemical Company (10005583) | 10 µM |
| | | Retinoic Acid (Freshly added) | Sigma-Aldrich (R2625-50MG) | 0.05 µM |
| | | FBS | Fisher Scientific Sweden (11573397) | 5 % |
| | Proximal differentiation medium | Proximal B | *Described above* | - |
| | | L-Glutamine | Fisher Scientific Sweden (12569079) | 1.5 mM |
| | | ITS | Fisher Scientific Sweden (12097549) | 1X |
| | | Cholera Toxin | Sigma-Aldrich (C8052-.5MG) | 0.1 µg/ml |
| | | Murine EGF | PeproTech (315-09-100ug) | 0.025 µg/ml |
| | | BPE | Fisher Scientific Sweden (11568866) | 30 µg/ml |
| | | Retinoic Acid (freshly added | Sigma-Aldrich (R2625-50MG) | 0.05 µM |
| | | BSA | Saveen & Werner (B2000-500) | 0.1 % |
| | Proximal organoid medium | Proximal B | *Described above* | - |
| | | ITS | Fisher Scientific Sweden (12097549) | 1X |
| | | Murine EGF | PeproTech (315-09-100ug) | 0.025 µg/ml |
| | | Cholera Toxin | Sigma-Aldrich (C8052-.5MG) | 0.1 µg/ml |
| | | BPE | Fisher Scientific Sweden (11568866) | 30 µg/ml |
| | | Retinoic Acid (freshly added) | Sigma-Aldrich (R2625-50MG) | 0.01 µM |
| | | FBS | Fisher Scientific Sweden (11573397) | 5 % |

**Table S2: Top genes per cell type used in deconvolution. Related to figure S3 and figure 4.** Gene lists are obtained using the function FindAllMarkers() on Seurat in GSE141259 Genes with logFC of higher than 0.5 displayed in this table.

| Cell type | Genes |
|---|---|
| ATI cells | Ager, Akap5, Anxa1, Anxa3, App, Aqp5, Cldn18, Clic3, Clic5, Clu, Cryab, Ctnna1, Ctsh, Emp2, Icam1, Igfbp7, Itgb6, Lgals3, Msn, Myl12a, Myl6, Ndnf, Phldb2, Serpinb6b, Serpinb9, Sparc, Timp3, Tpm1, Tpm3, Vamp8, Vegfa, |
| ATII cells | Abca3, Acot7, Acsl4, Atp5e, Atrx, Cd36, Ctsc, Cxcl15, Dbi, Dram1, Egfl6, Elovl1, Fabp5, Fasn, H2-K1, Hc, Ifi27l2a, Il33, Lamp3, Lcn2, Lpcat1, Lrg1, Lrp2, Ly6e, Lyz1, Lyz2, Malat1, Napsa, Npc2, Ptgs1, Rgcc, Rnase4, S100g, Scd1, Sfta2, Sftpa1, Sftpb, Sftpc, Sftpd, Slc34a2, Slpi, Tgoln1, |
| Basal Cells | Abi3bp, Apoe, Aqp3, Aqp4, Bpifa1, Dapl1, Dcn, Dst, Eef1a1, Eef1b2, Egr1, Ehf, Emb, Epas1, F3, Fxyd3, Glul, Gsta4, Gstm2, Gsto1, Jun, Krt15, Krt19, Krt5, Ltf, mt-Co1, Pabpc1, Rpl13a, Rpl14, Rpl18a, Rpl21, Rpl22, Rpl28, Rpl3, Rpl31, Rpl37a, Rpl5, Rpl8, Rplp2, Rps12, Rps16, Rps2, Rps3, Rps4x, Rps8, Rps9, Rpsa, S100a6, Tgm2, Tmem176a, Tmem176b, Tpt1, Zfp36l1, |
| Ciliated Cells | Acsl3, Aebp1, Agr3, Ak7, Akap14, Akap9, Aldh1a1, Aldh3b1, Anxa2, Anxa8, Aoc1, Arhgap18, Arl3, Ascc1, Atpif1, AU040972, B9d1, BC051019, Bphl, Calm1, Calml4, Capsl, Ccdc113, Ccdc146, Ccdc153, Ccdc17, Ccdc173, Ccdc181, Ccdc30, Ccdc39, Ccdc78, Cd24a, Cd55, Cdhr3, Cdhr4, Cep290, Cetn2, Cetn4, Chchd10, Chchd2, Chchd6, Ckb, Crip2, Cspp1, Csrp2, Cxcl17, Cyp2s1, Cyp4b1, Ddx5, Dnah12, Dnah3, Dnah5, Dnah6, Dnali1, Dpy30, Drc1, Dusp14, Dynll1, Dynlrb2, Efcab1, Efcab10, Eif1, Elof1, Enkur, Erich2, Ezr, Fam161a, Fam183b, Fam216b, Fhad1, Foxj1, Fth1, Gm1673, Gm973, Gsn, Gstm1, H3f3b, Hes6, Hint1, Hsp90aa1, Hspa4l, Hsph1, Ifitm1, Ift43, Ift74, Ift81, Ift88, Igfbp5, Iqca, Iqcg, Itm2b, Kif21a, Lgals3bp, Lrrc23, Lrrc51, Lrrc6, Lrrc71, Lrriq1, Map1b, Mcee, Meig1, Mlf1, Mns1, Mrps17, Mt1, Mycbp, Nek5, Nme5, Nudc, Nudt4, Odf3b, Osbpl6, Pcp4l1, Pebp1, Pifo, Plcb3, Pltp, Ppia, Ppil6, Ppp1r36, Prdx5, Ptges3, Rfk, Riiad1, Rsph1, Rsph4a, Rsph9, S100a1, S100a11, Sec14l3, Slc25a5, Smim5, Sntn, Spa17, Spag17, Spag6, Spef2, Srsf10, Stk33, Stmnd1, Tcea3, Tekt1, Tm4sf1, Tmem107, Tmem212, Tomm7, Tppp3, Traf3ip1, Trp53bp2, Tspan1, Tuba1a, Tubb4b, Txnip, Vpreb3, Wdr66, |
| Club cells | Alas1, AW112010, B430010I23Rik, Bpifb1, Bst2, C1qb, Cbr2, Ccl17, Ccl6, Cd52, Cd74, Cldn10, Cp, Crip1, Cst3, Ctss, Cyp2f2, Fau, Fmo2, Fmo3, Gsta3, H2-Aa, H2-Ab1, H2-Eb1, Ifit1, Ifit2, Ifit3, Ifitm3, Iigp1, Isg15, Mgst1, mt-Cytb, mt-Nd2, Phf11d, Pigr, Prdx6, Psap, Reg3g, Rsad2, S100a4, Scgb1a1, Scgb3a1, Selenbp1, Spp1, Stat1, Tff2, Tmsb4x, Trf, Vim, Wfdc2, |
| Goblet cells | Adh7, Agr2, Chad, Gp2, Hp, Lypd2, Muc5b, Qsox1, Retnla, Scgb3a2, Sult1d1, |
| Krt8+ ADI | Anxa5, Areg, Arpc2, B2m, Calm2, Ccng1, Cd81, Cdkn1a, Cldn4, Cstb, Cystm1, Dstn, Epcam, Esd, Ftl1, Gas5, Gdf15, H2-D1, Hbegf, Hspb1, Ifrd1, Itgb1, Krt18, Krt7, Krt8, Lmo7, Ly6a, Mt2, Myl12b, Nupr1, Pkm, Rpl11, Rpl12, Rpl13, Rpl15, Rpl17, Rpl19, Rpl23, Rpl23a, Rpl26, Rpl27, Rpl27a, Rpl32, Rpl34, Rpl35a, Rpl37, Rpl38, Rpl4, Rpl6, Rpl9, Rplp0, Rplp1, Rps11, Rps13, Rps14, Rps15, Rps15a, Rps17, Rps18, Rps19, Rps20, Rps23, Rps24, Rps25, Rps26, Rps27, Rps27a, Rps27l, Rps29, Rps3a1, Rps5, Rps6, Rps7, S100a10, Sfn, Sprr1a, Sqstm1, Tnfrsf12a, Tnip3, Tpm4, Txn1, Ubb, |
| Mki67+ Proliferation | Actb, Anln, Anp32b, Arl6ip1, Birc5, Cenpe, Cenpf, Dek, Hmgb1, Hmgb2, Hmgn2, Hmmr, Hnrnpa2b1, Hnrnpa3, Hnrnpab, Hsp90b1, Kif23, Lig1, Mki67, Ncl, Npm1, Pcna, Prc1, Ptma, Ranbp1, Rpl35, Serbp1, Smc2, Smc4, Spc24, Stmn1, Top2a, Tpx2, Tuba1b, Tubb5, Ube2c, Ybx1, |
| Neuroendocrine Cells | Anp32a, Arl4a, Ascl1, Calca, Car8, Cbx5, Ccnd2, Cd9, Celf4, Chga, Chgb, Col8a1, Cplx2, Ddc, F5, Fos, Gadd45b, Gnas, Hsp90ab1, Hspa5, Id4, Igf2, Itm2c, Meg3, Meis2, Nktr, Nnat, Ntn4, Pam, Pcsk1, Peg3, Piezo2, Pnmal2, Ptn, Ptprn, Resp18, Rgs2, Scg2, Scg5, Spock3, Syt7, Tspan13, Ttc3, |

**Supplemental experimental procedures**

*Fused Deposition Modelling (FDM) 3D printing parameters*
Standard Tessellation Language (STL) models were sliced using Ultimaker Cura software (v4.0, Ultimaker) at a resolution of 0.1mm. Sliced models were printed using an Ultimaker 3 printer using PLA filaments (Creative Tools, www.creativetools.se). The 3D printer's nozzle diameter was 0.4 mm, infill printing temperature was set at 205 °C, and heated-bed temperature: 60 °C. Prints were sterilized by incubation in 70 % ethanol for a minimum of 1 hour.

*Preparation of the lung for classic isolation*
Lungs were prepared as previously described (Jansing et al., 2018). Briefly, distal dissociation solution (Table S1) was injected through the cannulated trachea followed by 300 µl of 1 % agarose (A9414, Sigma) followed by placing a surgical knot to close the trachea.

*Distal progenitor cell isolation*
Briefly, isolated murine lung lobes were incubated in distal dissociation solution (Table S1) for 45 minutes at RT. Lobes were then minced using forceps in Base/DNase medium (Table S1). Cells were subsequently filtered through 100, 40, and 10 µm meshes sequentially. Fibroblasts and macrophages were removed by allowing single cell suspensions to attach on the surface of 10cm tissue culture treated petri dishes (Sarstedt, Sweden) for 30 minutes at 37 °C (2 dishes per mouse). Macrophages, monocytes, and endothelial cells were further depleted using the Quadri MACs system (130-091-051, Miltenyi Biotec, Sweden) using a mixture of anti-CD45 (130-052-301, Miltenyi Biotec) and anti-CD31 (130-097-418, Miltenyi Biotec) microbeads. EpCAM+ (130-105-958, Miltenyi Biotec) Distal progenitor cells were selected for organoid formation assays or for cell pellets used in RNA sequencing.

*Distal organoid formation*
Briefly, CCL206 cells (ATCC), murine lung fibroblasts (Mlg), were treated with 10 µg/ml Mitomycin C (M4287-2MG, Sigma-Aldrich, Sweden) for 1 hour at 37 °C to inhibit proliferation, followed by incubation with culture medium for at least 2 hours before trypsinization. A 1:1 ratio of CCL206/Matrigel to EpCAM+ distal progenitor cells was pipetted into a 6.5 mm 0.4 µm Corning trans-wells (10482181, Fisher Scientific) and incubated with distal organoid culture medium (Table S1). 20,000 cells of each CCL206 and distal progenitors were used per transwell at a final volume of 100 µl. Distal organoid medium was changed every 48 hours for 14 days.

*Organoid Counting and size distribution*
Bright field images of all organoid cultures were captured using a wide-field camera in a Cytation5 Multimodal imager/plate reader with a 4X objective (AH Diagnostics). Each transwell culture was imaged as a mosaic with z-stacking of 3 planes across the transwell culture with a distance of 219 µm between each plane. All organoids were visible in each z-plane at different focal planes, however, a focal stacking of all 3 planes resulted in a focused 2D image for all organoids making it suitable for analysis. The selection tool in FIJI was used to quantify the number of individual organoids and their size distribution based on area.

*Air liquid interface (ALI) of proximal progenitors*
ALI cultures of proximal progenitors were done as previously described (Eenjes et al., 2018). Briefly, expanded cells from individual tracheas were seeded on 6.5 mm 0.4 µm transwell inserts (10482181, Fisher Scientific) and submerged in proximal progenitors' proliferation medium until confluence (typically day 7) (Table S1). Cells were lifted to air after reaching confluence by only placing liquid on the basal side of the transwell and proximal differentiation medium was changed every 48 hours for 28 days.

*Proximal organoid formation*
Proximal organoids were grown as previously described (Rock et al., 2009; You et al., 2002). Expanded cells from individual tracheas were seeded in 100 µl Matrigel (11543550, Fisher Scientific, Sweden) at 10 000 cells/insert and were incubated in MTEC organoid medium for 14 days with media changes every 48 hours.

*Scanning Electron Microscopy (SEM)*
Differentiated proximal lung epithelial cells in ALI were fixed in 2.5 % Glutaraldehyde (G5882, Sigma-Aldrich) in PBS at 4 °C overnight and washed in PBS. Fixed samples were dehydrated in an ethanol series 50, 70, 80, 90, 100, 100 % for 10 minutes each at RT while they remained in the insert to avoid warping of the ALI membrane. Critical point drying (CPD) was done using K850 critical point dryer (K80, Quorum, UK). Dried ALI membranes were separated from the inserts and mounted face-up on SEM mounts (PELCO Tabs™, Carbon Conductive Tabs, 25 mm OD, Tedpella) and were sputter coated with platinum/palladium 80/20 at a thickness of 10 nm. Images were acquired using a Jeol JSM-7800F SEM microscope.

*Histology, Immunohistochemistry (IHC), and Immunofluorescence (IF).*
For histological assessment, samples were fixed in 10 % neutral-buffered formalin at 4 °C overnight and were processed using a xylene-free approach, using graded ethanol series and isopropanol (both Fisher Scientific, UK) before paraffin embedding (Histolab, Sweden). 4 µm wide sections were sliced and placed on microscope slides (Thermo Scientific, Germany) for staining. Deparaffinized sections was followed by staining with hematoxylin and eosin (Merck Millipore, Germany) followed by dehydration in consecutively graded ethanol series. Dried sections were mounted with Pertex (Histolab, Sweden). Antigen retrieval was performed with 10 mM citrate buffer (pH 6.0) in a pressure cooker for 20 minutes at 105 °C. For IF of paraffin sections, blocking was done for 1 hour at RT using 1X PBS / 5 % goat serum / 0.3 % Triton™ X-100, primary antibodies (AB) were diluted in AB dilution buffer consisting of 1X PBS / 1 % BSA / 0.3 % Triton™ X-100 and incubated at 4 °C overnight. The sections were incubated with phalloidin (1:100, A12379, Life Technologies) and the secondary antibodies at RT for 1 hour followed by 4′,6-diamidino-2-phenylindole (DAPI) staining at 5 µg/ml for 10 minutes at RT.

*Light-sheet sample preparation and imaging of distal lung organoids*
Distal organoid cultures were fixed in 10 % formalin (neutral-buffered and methanol-free) for 2 hours at RT and washed in PBS. Sample preparation and immunolabeling were done using the iDISCO protocol with some modifications (Renier et al., 2014). First, samples were dehydrated in methanol / PBS series (20, 40, 60, 80, 100, 100 % methanol in PBS) for 15 minutes for each step at RT. Delipidation was performed by incubating the samples in a mixture of dichloromethane (DCM, Sigma-Aldrich 270997) and methanol (66 % DCM / 33 % methanol) for 3 hours at RT. Samples were then rehydrated using a methanol / PBS series in the opposite order for 15 minutes each at RT. Samples were then incubated in permeabilization solution (20 % DMSO / 0.16 % Triton-X100 / 2.3 % glycine in PBS) overnight at 37 °C followed by incubation in blocking solution (6 % goat serum / 10 % DMSO / 0.168 % Triton-X100 in PBS) overnight at 37 °C. Primary antibodies were incubated overnight at 4 °C followed by washing in PBS 3 times while shaking for 1 hour each time. Secondary antibodies were incubated for 2 hours at RT and DAPI was incubated for 15 minutes at RT followed by washing in PBS 3 times while shaking for 1 hour each time. After immunolabeling, organoid samples were taken out of the inserts and embedded in 1 % low-gelling temperature agarose (Sigma Aldrich A9414) in milliQ water followed by dehydration using the methanol / PBS series for 1 hour in each step at RT. Samples were then incubated in DCM / methanol (66 % / 33 %) for 1 hour followed by 2 changes of 100 % DCM for 15 minutes each while shaking. Reflective index (RI) matching of the processed samples was done by incubating the sample in ethyl cinnamate (RI: 1.56, Sigma-Aldrich, 112372-100G) until imaging. Imaging was done using the Aurora airy beam LSFM system equipped with two multi-immersion objectives (Special Optics 0.4 NA @ 1.45 RI (RI 1.33-1.56; Mag 15.3x-17.9x) WD 12 mm FOV 870nm 1 µum axial resolution) and Hamamatsu Orca-Flash 4.0 V3 digital CMOS camera (M Squared Lasers Ltd.). Maximum intensity projection or volumetric data of these images was obtained using either Arivis Vision4D or Imaris Viewer V9. Segmentation of each organoid was done using virtual reality (VR) software SyGlass.

*mRNA library preparation and sequencing and bioinformatics*
Library preparation was done using the Illumina Stranded mRNA Prep Ligation (20040534, Illumina) with 14 cycles under final PCR amplification and samples were indexed with IDT for Illumina RNA UD Indexes Set B, Ligation (20040554, Illumina) for multiplexing. Clean up steps were automated using the King Fisher FLEX (18-5400620, Thermo Scientific). Library concentration was checked using the QuantIT 1X dsDNA HS Assay Kit (Q33232) and library size and quality were checked using the 5200 Fragment Analyzer™ 12-capillary (M3510AA, Agilent) using the DNF-930 dsDNA Reagent Kit, 75 bp – 20,000 bp (500 Samples) (Part # DNF-930-K0500, Agilent). A library input of 1.25 nM was sequenced using NovaSeq 6000 Sequencing System (20012850, Illumina). Alignment was done using STAR align and read counts were extracted using featureCounts (Dobin et al., 2012) (Liao et al., 2013). All reference genomes were from the Ensembl database (Mouse GRCm38 with the GTF annotation (release 101). Analysis of RNAseq data was done on R using DEseq2, ggplot2, EnhancedVolcano, and pheatmap. Z-score is calculated based on substracting the mean TPM values of all samples in each gene from the TPM value for each sample for each gene divided by the standard deviation of the TPM values per gene across all samples.

$$\text{z-score: } \frac{Gene\ TPM_{(sample)} - Gene\ TPM_{(mean\ of\ all\ samples)}}{standard\ deviation}$$

**Supplemental references**


Dobin, A., Davis, C.A., Schlesinger, F., Drenkow, J., Zaleski, C., Jha, S., Batut, P., Chaisson, M., and Gingeras, T.R. (2012). STAR: ultrafast universal RNA-seq aligner. Bioinformatics *29*, 15-21. 10.1093/bioinformatics/bts635.

Eenjes, E., Mertens, T.C.J., Buscop-van Kempen, M.J., van Wijck, Y., Taube, C., Rottier, R.J., and Hiemstra, P.S. (2018). A novel method for expansion and differentiation of mouse tracheal epithelial cells in culture. Scientific Reports *8*, 7349. 10.1038/s41598-018-25799-6.

Jansing, N.L., McClendon, J., Kage, H., Sunohara, M., Alvarez, J.R., Borok, Z., and Zemans, R.L. (2018). Isolation of Rat and Mouse Alveolar Type II Epithelial Cells. In Lung Innate Immunity and Inflammation: Methods and Protocols, S. Alper, and W.J. Janssen, eds. (Springer New York), pp. 69-82. 10.1007/978-1-4939-8570-8_6.

Liao, Y., Smyth, G.K., and Shi, W. (2013). featureCounts: an efficient general purpose program for assigning sequence reads to genomic features. Bioinformatics *30*, 923-930. 10.1093/bioinformatics/btt656.

Renier, N., Wu, Z., Simon, David J., Yang, J., Ariel, P., and Tessier-Lavigne, M. (2014). iDISCO: A Simple, Rapid Method to Immunolabel Large Tissue Samples for Volume Imaging. Cell *159*, 896-910. https://doi.org/10.1016/j.cell.2014.10.010.

Rock, J.R., Onaitis, M.W., Rawlins, E.L., Lu, Y., Clark, C.P., Xue, Y., Randell, S.H., and Hogan, B.L.M. (2009). Basal cells as stem cells of the mouse trachea and human airway epithelium. *106*, 12771-12775. 10.1073/pnas.0906850106 %J Proceedings of the National Academy of Sciences.

You, Y., Richer, E.J., Huang, T., and Brody, S.L. (2002). Growth and differentiation of mouse tracheal epithelial cells: selection of a proliferative population. American journal of physiology. Lung cellular and molecular physiology *283*, L1315-1321. 10.1152/ajplung.00169.2002.
